# Selection for male aggression is associated with changes in reproductive traits, chemical signaling and lifespan in *Drosophila melanogaster*

**DOI:** 10.1101/2025.06.02.657383

**Authors:** Anthony Defert, Romane Gout, Gaelle Pennot, Fanny Jamme, Amandine Castex, Anissa Handjar, Thomas Guilleman, Jean-Christophe Billeter, Séverine Trannoy

**Affiliations:** Research Center on Animal Cognition (CRCA), Center for Integrative Biology, Toulouse University, CNRS, UPS, Toulouse, France; Institute for Zoology, Halle-Wittenberg University, Halle (Saale), Germany; Groningen Institute for Evolutionary Life Sciences, University of Groningen, 9747AG Groningen, The Netherlands

**Keywords:** Life history traits, trade-off, Aggression, Reproduction, Survival, chemical signaling, *Drosophila melanogaster*

## Abstract

Life history traits are all essential traits linked to development, survival and reproduction, directly influencing individual fitness. Limited environmental or physiological resources force organisms to balance competing demands, leading to fundamental trade-offs, particularly between survival and reproduction. Male aggression is often assumed to enhance reproductive success, yet its broader association with other fitness-related traits remains unclear. To address this, we used *Drosophila melanogaster Bully* lines selected for high levels of male aggression, to examine relationships between aggression, chemical signaling, and essential life-history traits including reproductive behaviors and lifespan. Our results reveal that *Bully A* males exhibit a shift in a major life-history trade-off, with reduced reproductive performance accompanied by increased lifespan. *Bully A* males exhibited lower mating success, shorter mating duration and less effective chemical mate-guarding. However, this was traded-off with increased lifespan, enabling more reproductive opportunities. We also uncover potential mechanistic underpinnings of the reproductive deficits. *Bully* males showed differences in their cuticular hydrocarbon (CHCs) profiles and transferred lower levels of cVA to females, a mate-guarding pheromone, thereby weakening their post-mating strategy. Overall, our findings indicate that selection for male aggression is associated with a shift in a key life-history trade-off, highlighting aggression as an evolutionary force shaping adaptation of life-history traits and providing a foundation for future genetic and mechanistic studies.

## INTRODUCTION

Life history traits, such as development time, fertility, survival, and reproductive success, are key phenotypic components that directly influence individual fitness (Braendle and Paaby, 2024; Roff, 1992; Stearns, 1992). These traits often face trade-offs: investing in one can compromise another (Fisher, 1930; Stearns, 1989; Williams, 1966). For instance, high reproductive effort may reduce survival due to the energetic costs of gamete production or increased mortality associated with mating behaviors or competition (Flatt, 2011; 2020). Trade-offs also emerge between pre- and post-copulatory strategies (Simmons, 2001; Warner et al., 1995). Males prioritizing pre-copulatory investment, such as aggressive competition for mates, may have limited resources for post-copulatory investment linked to sperm competition such as mate guarding (Andersson, 1994; Brennan and Orbach, 2021). Understanding how such trade-offs are shaped by natural selection offers insight into how organisms optimize fitness in competitive social environments (Roff, 1992; Stearns, 1976).

The fruit fly *Drosophila melanogaster (D. mel.)*, provides a powerful model for studying life-history trade-offs, with well-characterized developmental, behavioral, and reproductive traits (Flatt, 2020). These traits are shaped by both genetic and environmental factors and can evolve under ecological pressures. Faster development, earlier reproduction and high mating rate, for instance, are advantageous in changing environments but come at the cost of reduced longevity and fecundity (Partridge and Farquhar, 1981; Stearns, 2000). Experimental manipulations of environmental conditions such as temperature and food availability reveal how external factors shape trade-offs between survival and reproduction (Chippindale et al., 1996; Marshall and Sinclair, 2010).

Beyond development and fecundity, *D. mel.* serves as a model to study a wide range of social behaviors, including aggregation, aggression, and courtship, that are largely mediated by chemical communication (Kim et al., 2017; Kohl et al., 2015). Cuticular hydrocarbons (CHCs) such as some pheromones play diverse roles in social interaction: 7-Tricosene (7-T), a male-specific compound, increases female receptivity and modulates aggression (Grillet et al., 2006; Wang et al., 2011); cis-vaccenyl acetate (cVA), a male-transferred pheromone, attracts females, repels other males, and promotes aggregation (Kurtovic et al., 2007; Laturney and Billeter, 2016); and the female-specific hydrocarbon 7,11-heptacosadiene (7,11-HD) strongly elicits male courtship and supports species-specific mate recognition (Toda et al., 2012).

Among these interactions, male-male competition plays a central role in shaping reproductive strategies. Pre-copulatory competition involves resource defense, courtship displays, and aggressive interactions to secure mating opportunities. Courtship in *D. mel.* includes stereotyped behaviors such as orientation, wing extensions (producing courtship song), tapping, licking, and attempted copulation (Yamamoto and Koganezawa, 2013). Successful courtship displays lead to female acceptance, outcompeting other males that fail to meet the female’s preferences (Baxter et al., 2018; Hindmarsh Sten et al., 2025b). Aggression intensifies this competition: more aggressive or dominant males often monopolize access to females (Dow and von Schilcher, 1975; Filice and Dukas, 2019; Gao et al., 2024; Nandy et al., 2016; Prunier and Trannoy, 2024). Aggressive behaviors range from wing threats and lunging to high-intensity physical confrontations like tussling and boxing (Chen et al., 2002). Beyond aggressive patterns, males also produce agonistic wing flicks when competing for access to females, which can interfere with a female’s auditory perception of rival males and thereby influence mate choice (Hindmarsh Sten et al., 2025a). These behaviors promote territorial dominance and enhance mating success (Gao et al., 2024), although subordinate males may also achieve reproductive success through first access to females and prolonged copulation (Filice and Dukas, 2019).

Post-copulatory competition further shapes male reproductive success. After mating, males use multiple strategies to ensure paternity, including the transfer of seminal fluid proteins (SFPs) including Sex Peptide (SP) and Accessory gland proteins (Acps), which promote sperm storage, stimulate oviposition, and reduce female receptivity to further mating (Chapman et al., 2003; Fricke et al., 2009). These factors also contribute to the formation of a mating plug that retains sperm, acting as a passive mate-guarding strategy (Avila et al., 2011; Dunham and Rudolf, 2009). In parallel, chemical mate guarding via pheromone transfer (e.g. cVA and 7-T) reduce female attractiveness and their likelihood of re-mating (Billeter et al., 2009; Gillott, 2003; Laturney and Billeter, 2016; Miyamoto and Amrein, 2008; Scott, 1986; Zawistowski and Richmond, 1986). Male reproductive behavior is plastic: in the presence of rivals, males extend their mating duration (Bretman et al., 2009; Bretman et al., 2013; Kim et al., 2013), increase the number of sperm in their ejaculate (Garbaczewska et al., 2013), adapt seminal fluid content in their ejaculate (Hopkins et al., 2019), and present higher levels of aggression toward opponents to guard their mates (Baxter et al., 2015; Bretman et al., 2011; Dore et al., 2021; Mazzi et al., 2009). Together, this demonstrates that male-male aggression influences access to mates and reproductive outcomes through both pre- and post-copulatory competition.

While aggression can enhance male mating success in natural settings, the relationship between aggression and reproductive fitness is not always straightforward. In a previous study, Penn et al. (Penn et al., 2010) generated “*Bully*” flies through 37 generations of selection for aggression starting from a *Canton-S* (*Cs*) population. In the first step, *Cs* winner males from aggressive encounters were allowed to mate with *Cs* females. In the resulting F1 progeny, males displaying higher levels of aggression were again selected and mated with their female siblings. The resulted hyper-aggressive males frequently escalated to boxing, the most intense aggression phenotype (Chowdhury et al., 2017), but showed reduced reproductive success with virgin females compared to unselected control males (Penn et al., 2010). This suggests that although aggression may enhance competitive ability, it can also incur fitness costs. This raises important questions about the role of aggression in shaping life-history trade-offs, such as: to what extent does aggression influence life history traits related to reproduction, and does it affect the balance reproduction vs survival?

To address these questions, we investigated whether *D. mel.* males from *Bully* lines exhibit changes in life-history trade-offs compared to the unselected starting wild-type flies (*Cs* lines). We found that hyper-aggressive *Bully* males show reduced reproductive success and shorter mating duration across their lifespan, indicating lower pre-copulatory success. Their mates present higher remating rate and increased attractiveness, suggesting that mate-guarding strategy of *Bully* males is less efficient. However, this poor pre- and post-copulatory success was traded-off with an increased lifespan, which is accompanied by late-life reproductive success. We then provided a mechanistic basis for the reduced copulatory success of *Bully* males. They showed distinct CHCs profiles compared to *Cs* males, which may contribute to their lower mating success. Moreover, they transferred to females lower levels of cVA, a mate-guarding pheromone, to females, potentially decreasing efficiency of their post-mating strategy. Our findings underscore the complex interplay between aggression, reproductive behavior, survival, and chemical signaling, providing insights into the evolutionary dynamics of life-history trade-offs in competitive social contexts.

## RESULTS

### Males exhibiting higher aggression intensity also exhibit shorter mating durations

To determine whether selection for male aggressive traits may also be associated with changes in reproductive behaviors, we employed three male lines selected for increased aggression. In a previous study, Penn et al. (Penn et al., 2010) generated the so-called “*Bully*” flies through 37 generations of artificial selection for male aggression starting from a *Canton-S* (*Cs*) population. Briefly, males that won aggressive encounters were allowed to mate with *Cs* females. In the resulting F1 progeny, males exhibiting higher levels of aggression were again selected and mated with their female siblings. Following the same experimental procedure, several independent selections were performed in parallel, giving rise to the *Bully A, Bully B and Bully C* lines. Although the *Bully* lines originated from a *Canton-S* population, they subsequently evolved independently during the artificial selection procedure and continued to be maintained as separate laboratory populations. Consequently, some phenotypic differences may reflect genetic divergence accumulated during independent maintenance, in addition to the effects of selection for aggression. Males from these three lines (*Bully* A, B, and C) were tested in standardized aggression assays (Trannoy et al., 2015b) (Figure 1A). Across same-line contests, males from all three *Bully* lines did not differ from *Cs* males in latency to lunge (Figure 1B), but displayed significant increase in the number of lunges (Figure 1C), the number of boxing events (Figure 1D), and boxing frequency (defined as the proportion of male pairs displaying at least one boxing event during the 10-min observation period) (Figure 1E) compared to *Cs* males. However, the magnitude of this phenotype differed across lines: *Bully A* and *Bully B* males exhibited higher aggression levels than *Bully C* males (Figure 1E).

**Figure 1.**
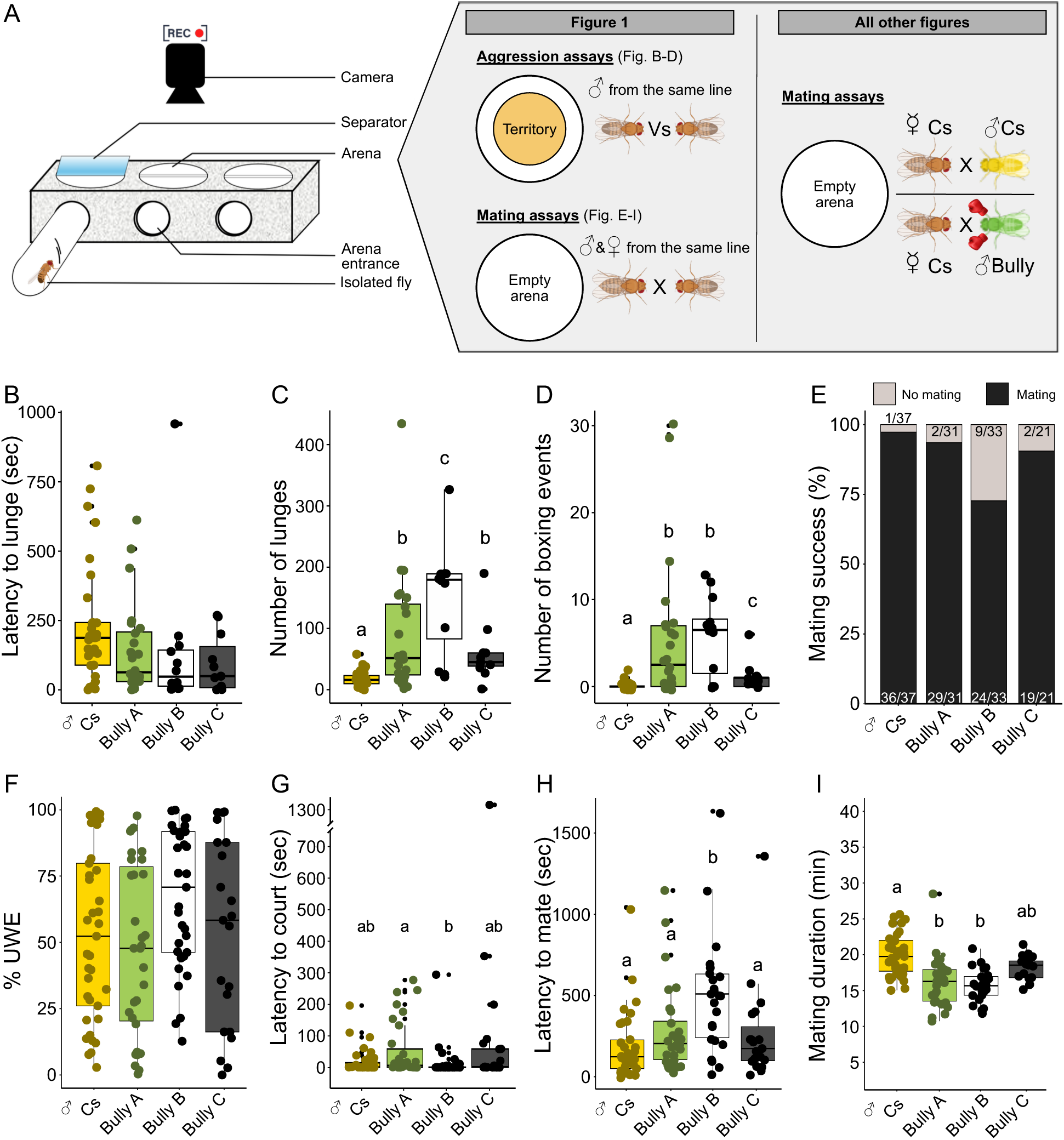
Courtship and aggressive behaviors in *Cs*, *Bully* A, *Bully* B and *Bully* C lines. (A) Schema of the experimental set up to record aggressive and courtship behaviors. Couples of flies and presence of food cup are mentioned according to the figure of this article. (B) Latency to lunge of *Cs*, *Bully A*, *Bully B* and *Bully C* males in aggression assay with a territory against a male of the same line (n*_Cs_*=29, n*_BullyA_*=24, n*_BullyB_*=12, n*_BullyC_*=11, LM and permutation test (Monte-Carlo test): P=0.136). (C) Number of lunges of *Cs*, *Bully A*, *Bully B* and *Bully C* male pairs in aggression assays (n*_Cs_*=29, n*_BullyA_*=24, n*_BullyB_*=12, n*_BullyC_*=11, GLM quasi-Poisson and type II analysis of deviance table (Likelihood Ratio Test): P=1.622×10^−10^, LRχ²=48.556, post-hoc details in Figure 1-table supplement 1). (D) Number of boxing events of *Cs*, *Bully A*, *Bully B* and *Bully C* male pairs in aggression assays (n*_Cs_*=29, n*_BullyA_*=24, n*_BullyB_*=12, n*_BullyC_*=11, GLM Poisson and type II analysis of deviance table (Likelihood Ratio Test): P=2.2×10^−16^, LRχ²=2511.9). (E) Non-competitive mating success of *Cs*, *Bully A*, *Bully B* and *Bully C* males paired with a virgin *Cs* female (n*_Cs_*=37, n*_BullyA_*=31, n*_BullyB_*=33, n*_BullyC_*=21, GLM binomial and type II analysis of deviance table (Likelihood Ratio Test): P=0.012, LRχ²=11.040). (F) Proportion of time spent by *Cs* and *Cs*, *Bully A*, *Bully B* and *Bully C* males doing Unilateral Wing Extension (UWE) toward a *Cs* virgin female in non-competitive courtship assays (n*_Cs_*=37, n*_BullyA_*=31, n*_BullyB_*=33, n*_BullyC_*=21, LM and type II Anova table: P=0.088, F=2.235). (G) Latency to court (time difference between first interaction and first UWE) of *Cs*, *Bully A*, *Bully B* and *Bully C* males paired with a virgin *Cs* female (n*_Cs_*=37, n*_BullyA_*=31, n*_BullyB_*=33, n*_BullyC_*=21, LM and type II Anova table: P=0.014, F=3.707). (H) Latency to mate (time difference between first interaction and start of the mating) of *Cs*, *Bully A*, *Bully B* and *Bully C* males paired with a virgin *Cs* female (n*_Cs_*=36, n*_BullyA_*=29, n*_BullyB_*=24, n*_BullyC_*=19, LM and type II Anova table: P=8.527×10^−4^, F=5.970). (I) Mating duration of *Cs*, *Bully* A, *Bully B* and *Bully C* males paired with a virgin *Cs* female (n*_Cs_*=36, n*_BullyA_*=29, n*_BullyB_*=24, n*_BullyC_*=19, LM and type II Anova table: P=5.015×10-6, F=10.365). For all graphs, different letters mean a statistical difference. Post-hoc analyses and statistical details are given in Figure 1-table supplement 1.

We next assessed courtship and mating behaviors by pairing males and females from the same line and scored courtship metrics. The proportion of time spent by males displaying unilateral wing extension (UWE) and latency to court did not differ significantly between lines (Figure 1F-G), while *Bully B* males exhibited a longer latency to mate compared to other pairings (Figure 1H). In contrast, mating duration (MD) was significantly reduced in *Bully A* and *Bully B* pairings compared to *Cs* (Figure 1I), while *Bully C* males again show an intermediate phenotype (Figure 1I).

These results indicate that males from the *Bully A* and *Bully B* lines, classified as hyper-aggressive in same-line competition (Figure 1B-E), also exhibit reduced mating duration when paired with same-line females, whereas *Bully C* males do not (Figure 1I). Because the *Bully A* line (hereafter named *Bully*) has been already employed in previous studies to measure differential gene expression profile (Chowdhury et al., 2017; Penn et al., 2010), we chose to conduct subsequent experiments on this line. Overall, these results support our initial hypothesis that selection for aggressive traits may have influenced other behavioral traits linked to reproductive behaviors.

### Highly aggressive males exhibit reduced mating success and have shorter mating duration

As the selection for aggression was male-specific (Penn et al., 2010), we next aimed to focus on examining male reproductive behaviors while avoiding confounding effects of female behavioral variation. Thus, all subsequent courtship and mating assays were performed by pairing *Cs* or *Bully A* males with *Cs* females, which served as a standardized female background. We first assessed mating success in competitive assays, where a *Cs* male and a *Bully* male competed for access to a single female. Under these conditions, no significant mating advantage was observed for *Bully* males (Figure 2A). We then assessed mating success in non-competitive assays by pairing each male with a virgin *Cs* female for 30 minutes. Under these conditions, *Bully* males showed a significantly reduced probability of mating compared to *Cs* males (Figure 2B), despite similar levels of courtship behavior, as measured by UWE (Figure 2C), courtship latency (Figure 2D), and mating latency among successful pairs (Figures 2E). We next examined copulation *per se* and found that MD was significantly shorter in *Bully* males, with a reduction of approximately 25% compared to *Cs* males (Figure 2F), consistent with the phenotype observed when males were paired with females from the same line (Figure 1I). Importantly, this reduction in MD persisted when *Bully* males were paired with decapitated *Cs* females (Figure 2G), indicating that a substantial contribution of the males and that active female behavioral responses controlled by the brain are unlikely to account for the observed reduction in mating duration.

**Figure 2.**
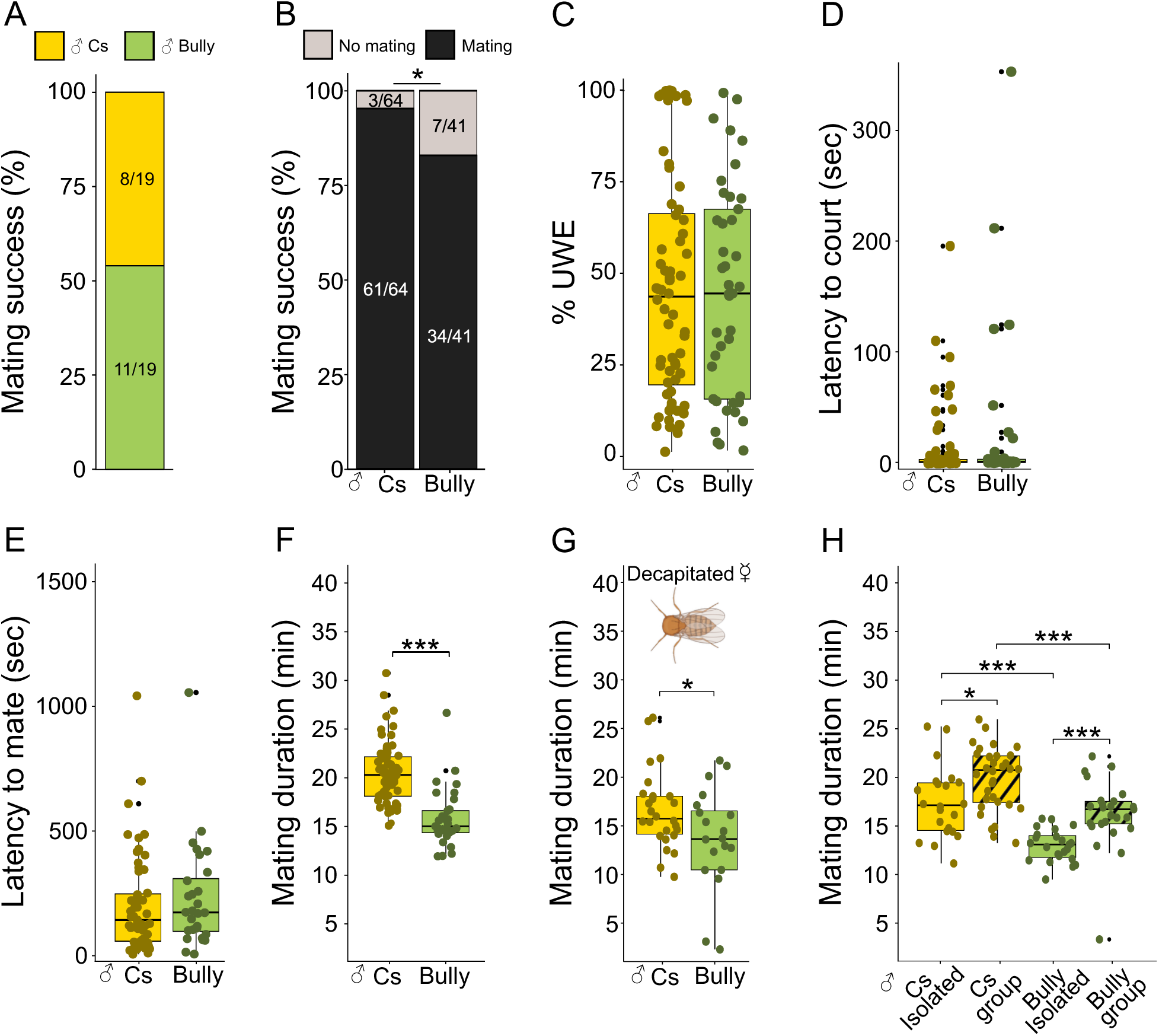
***Bully* males exhibit reduced mating success and allocate less time to mating.** (A) In competitive courtship assays, the mating success of *Cs* and *Bully* males was scored when paired with a single virgin *Cs* female (n*_Cs_*=19, n*_Bully_*=19, GLM binomial and type II analysis of deviance table (Likelihood Ratio): Test, P=0.329, LRχ²=0.951). (B) In non-competitive courtship assays, the mating success of *Cs* and *Bully* males paired with a virgin *Cs* female were analyzed (n*_Cs_*=64, n*_Bully_*=41, GLM binomial and type II analysis of deviance table (Likelihood Ratio Test): P=0.037, LRχ²=4.347). (C) Proportion of time spent by *Cs* and *Bully* males doing Unilateral Wing Extension (UWE) toward a *Cs* virgin female in non-competitive courtship assays (n*_Cs_*=64, n*_Bully_*=41, Fisher Pitman permutation test: P=0.877, t= −0.131). (D) Latency to court (time difference between first interaction and first UWE) of *Cs* and *Bully* males paired with a virgin *Cs* female (n*_Cs_*=64, n*_Bully_*=41, Fisher Pitman permutation test: P=0.319, Z=1.043). (E) Latency to mate (time difference between first interaction and start of the mating) of *Cs* and *Bully* males paired with a virgin *Cs* female (n*_Cs_*=61, n*_Bully_*=34, Fisher Pitman permutation test: P=0.266, Z=1.170). (F) Mating duration of *Cs* and *Bully* males paired with a virgin *Cs* female (n*_Cs_*=61, n*_Bully_*=34, T-test: P=6.009×10^−10^, t= −6.917) or (G) a virgin *Cs* decapitated female (n*_Cs_*=26, n*_Bully_*=20, T-test: P=0.022, t= −2.367). (H) Mating duration of *Cs* and *Bully* males that were previously raised in group of 10 males or in social isolation (n*_Cs-isolated_*=22, n*_Cs-group_*=32, n*_Bully-isolated_*=23, n*_Bully-group_*=25, LM and type II ANOVA: F-test, P=7.582×10^−11^, F=21.669). For all graphs, stars indicate significant differences (* P<0.05, ** P< 0.01, *** P<0.001). Post-hoc analyses and statistical details are given in Figure 1-table supplement 1.

As group-housed males have lengthened MD (Bretman et al., 2013; Kim et al., 2013), we compared MD of singly and group-housed males. We found that both *Cs* and *Bully* group-housed males exhibited significantly extended mating durations compared to single-housed males (Figure 1H). However, the MD in *Bully* males remained shorter compared to *Cs* males in group-housed condition (Figure 2H), indicating that *Cs* and *Bully* males respond equally to early social experience

Together, these results show that *Bully* males display reduced mating success and shorter copulation duration, indicating differences in pre-copulatory outcomes and copulatory investment. This supports our main hypothesis that selecting males for aggressive traits may also had influenced one or both pre- and post-copulatory strategies.

### Post-mating behaviors are affected in hyper-aggressive males

Given these differences in mating performance, we asked whether they compensate through distinct post-mating strategies. To investigate this, we first quantified courtship behavior immediately after copulation by measuring unilateral wing extension (UWE) directed toward the just-mated female. While *Cs* males performed minimal UWE towards just-mated females (Figure 3A’), *Bully* males displayed significantly higher levels of post-mating courtship in this context (Figure 3A’). This effect persisted when females were swapped after mating, with *Bully* males showing increased UWE towards *Cs*-mated females (Figure 3B–B’). This indicates that this phenotype reflects an intrinsic increase in courtship drive rather than a female-dependent effect. This elevated post-mating motivation led us to hypothesize that *Bully* males might achieve higher remating rates. To test this, we conducted sequential mating assays in which a single *Cs* or *Bully* male was consecutively paired with six virgin females over a 3-hour period and we scored the number of mating. Contrary to our expectation, *Bully* males did not mate with more females than *Cs* controls (Figure 3C), despite maintaining consistently shorter MD across successive mating (Figures 3D). As mating accompanies ejaculate transfer and incurs a reduction in sperm and seminal fluid reserve in *D. mel.* males (Hihara, 1981; Sirot et al., 2009), we further examined whether shorter MD in *Bully* males could reflect more efficient allocation, allowing fertilization of a greater number of females. However, analysis of progeny production revealed no significant differences between *Bully*- and *Cs*-mated females across successive matings (Figure 3E). Thus, increased post-mating courtship in *Bully* males does not translate into improved reproductive output.

**Figure 3.**
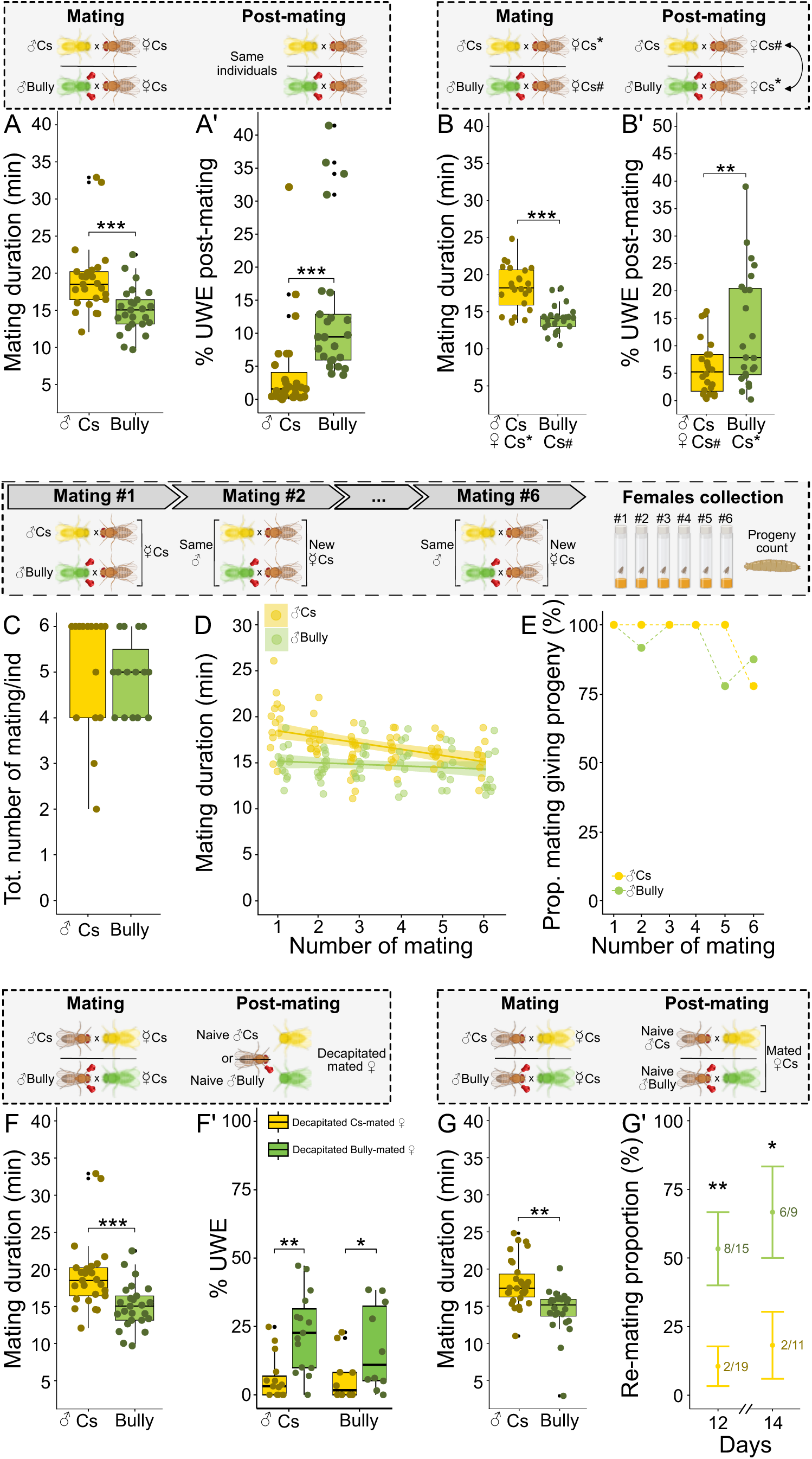
Hyper-aggressive males exhibit changes in mate-guarding features. (A, B, F, G). Mating duration of *Cs* and *Bully* males paired with a virgin *Cs* female (A: n*_Cs_*=21, n*_Bully_*=22, T-test: P=2.912×10^−11^, t= −8.759, B: n*_Cs_*=24, n*_Bully_*=24, T-test: P=5.425×10^−7^, t= −5.818, F: n*_Cs_*=25, n*_Bully_*=25, T-test: P=7.139×10^−4^, t= −3.617, G: n*_Cs_*=30, n*_Bully_*=24, T-test: P=8.743×10^−5^, t= −4.255). (A’) After mating ended, the proportion of time spent by each male to court performing UWE towards the just mated female was scored for 10 minutes (n*_Cs_*=21, n*_Bully_*=22, Fisher Pitman permutation test: P<0.001, Z=3.239). (B’) The proportion of time spent by the males performing UWE an unfamiliar mated female was scored for 10 minutes (n*_Cs_*=24, n*_Bully_*=24, Fisher Pitman permutation test: P=0.007, Z= −2.679). (C) Total number of mating reached by *Cs* and *Bully* males over six consecutive mating with virgin *Cs* females (type II analysis of deviance table (n*_Cs_*=18♂, n*_Bully_*=18♂, Wald Chi-square test): P=0.659, χ²=0.195). (D) Mating duration of *Cs* and *Bully* males over six consecutive mating with virgin *Cs* females (n*_Cs_*=18♂, n*_Bully_*=18♂, LMM and Satterthwaite T-test: P_interaction_=0.001, t= −3.261). A significant interaction between males’ genotype and number of mating was found, indicating that *Cs* decreased their mating duration with the number of mating, while *Bully* males did not. (E) Proportion of mating that gave rise to progeny in consecutive mating assay (n*_Cs_*=18♂, n*_Bully_*=18♂, GLMM and Wald Chi-square test: P_male_genotype_=0.279, χ²=1.173, P_mating_=0.847, χ²=2.019). (F’) Proportion of time spent by naïve *Cs* and *Bully* males to display UWE either toward *Cs*- mated or *Bully*-mated decapitated female over an observation period of 2 minutes (n*_Cs-Cs-mated_*=12, n*_Cs-Bully-mated_*=15, n*_Bully-Cs-mated_*=9, n*_Bully-Bully-mated_*=10, LMM and permutation test (Monte-Carlo test): P_male_genotype_<0.001, P_mated_female_=0.457). (G’) Proportion of *Cs*-mated and *Bully*-mated females that remated 12 or 14 days after a first mating (n*_Cs-12days_*=19, n*_Bully-12days_*=15, n*_Cs-14days_*=11, n*_Bully-14days_*=9, GLM binomial and type II analysis of deviance table (Likelihood Ratio Test): P_mated_female_=3.639×10^−4^, LRχ²=12.708, P_day_=0.385, LRχ²=0.756). For graphs A, A’, B, B’, F, F’, G and G’, stars indicate significant differences (* P < 0.05, ** P < 0.01, *** P < 0.001). Statistical details are given in Figure 1-table supplement 1.

We next investigated whether *Bully* males differ in their ability to modify female post-mating state. During mating, males transfer pheromonal cues, including cuticular hydrocarbons (CHCs), that reduce female attractiveness to rival males. We reasoned that the shorter MD of *Bully* males might limit the transfer of such mate-guarding signals, thereby reducing the extent of female unattractiveness. To test this independently of female behavior, we performed binary choice assays using decapitated females previously mated with either *Cs* or *Bull*y males. Both *Cs* and *Bully* males spent significantly more time courting females that had mated with *Bully* males (Figure 3F’, green vs yellow bars), indicating that these females remain more attractive and suggesting reduced transfer of inhibitory cues. To validate the experimental procedure and rule out potential deficits in sensory perception, we confirmed that both *Cs* and *Bully* males spent significantly more time performing UWE toward a decapitated female than toward a decapitated male (Figure 3-figure supplement 1). Finally, we assessed the functional consequences of this change in female state on remating behavior. Virgin *Cs* females were first mated with either *Cs* or *Bully* males and subsequently paired with naïve *Cs* males several days later. Females previously mated with *Bully* males, remated significantly more than females that had mated with *Cs* males (Figure 3G’, green), consistent with weaker mate-guarding.

Together, these results indicate that *Bully* males fail to offset their poor pre-copulatory success with more efficient post-mating strategies. Despite increased post-mating courtship motivation, they do not achieve higher remating rates or reproductive output, and their mates remain more attractive and more likely to remate. This dissociation indicates that increased courtship motivation is not predictive of reproductive success in this context. These findings suggest that selection for aggressive traits in males appears to compromise both pre- and post-copulatory aspects of male reproductive performance.

### Hyper-aggressive males present differences in their CHCs profiles and transfer lower quantities of mate-guarding pheromones to females

To investigate mechanisms underlying reduced mating success (Figure 2B) and changes in mate-guarding behaviors (Figure 3F-G) in *Bully* males, we quantified the CHCs profiles of naïve *Cs* and *Bully* males. PCA analysis revealed a clear separation between the two groups, indicating distinct CHCs profiles between naïve *Cs* and *Bully* males (Figure 4A). Quantification of individual compounds showed that *Bully* males exhibited significantly higher levels of alkanes, monoenes, and methyl-alkanes (Figure 4B, Table 1).

**Figure 4.**
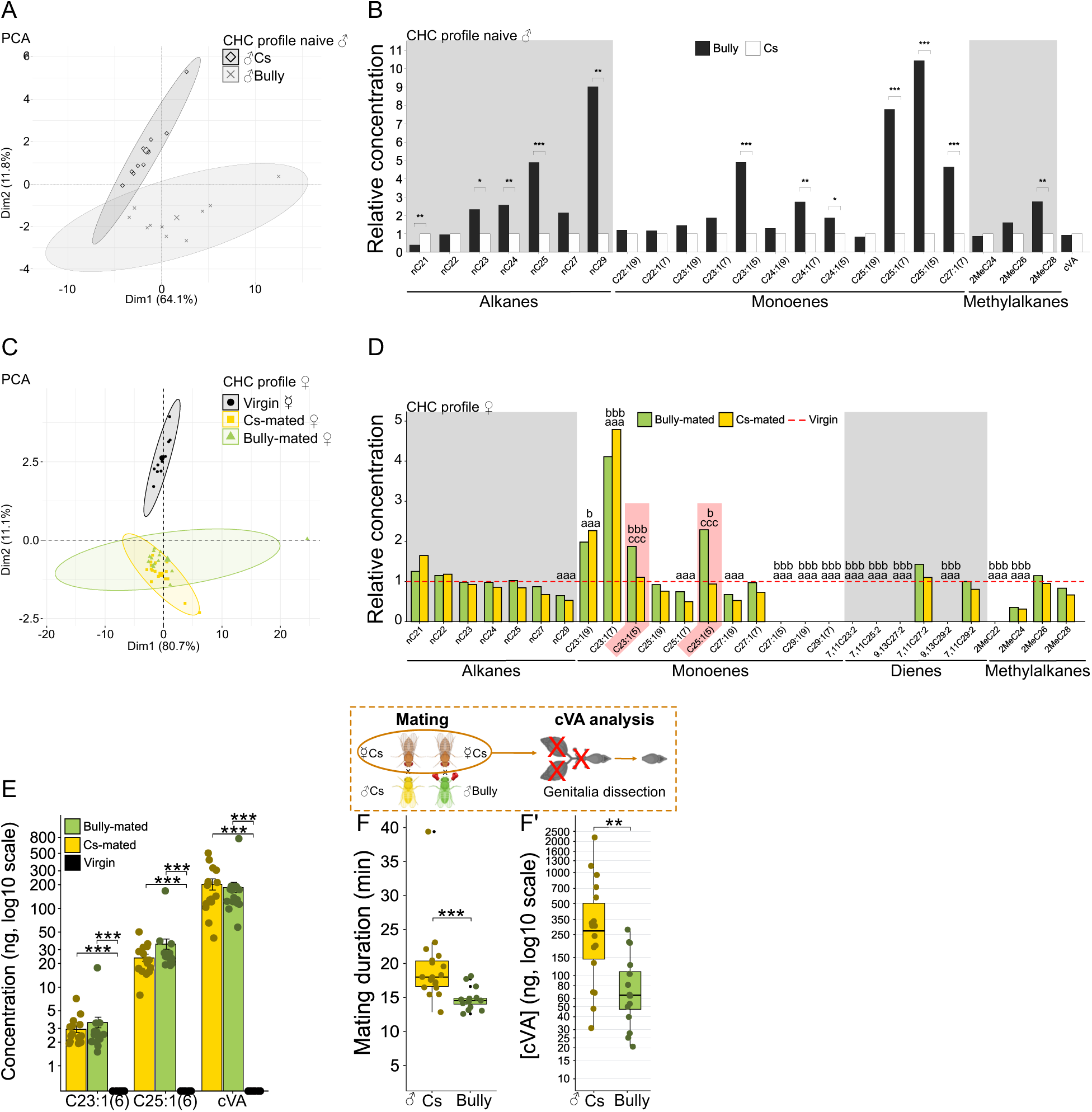
Hyper-aggressive males exhibit altered CHCs profiles and transfer reduced amounts of mate-guarding pheromones to females. (A) PCA analysis on mean concentration for each Cuticular Hydrocarbon (CHCs) compound of naïve *Cs* and *Bully*. Their CHCs profiles were found similar (n*_Cs_*=10♂, n*_Bully_*=10♂, PerMANOVA: P=0.002, F=6.789, R²=0.274). (B) Relative concentrations of *Bully*’s CHCs were calculated with respect to those of *Cs* males. The CHCs levels in *Cs* males were normalized to 1, and *Bully*’s CHC concentrations were expressed relative to this reference (n*_Cs_*=10♂, n*_Bully_*=10♂, LMM and permutation test (Monte-Carlo test): P=0.009). (C) PCA analysis on mean concentration for each CHCs compound of virgin, *Cs*-mated and *Bully*-mated females (n*_virgin_*=10♀, n_Cs-mated_=16♀, n*_Bully-mated_*=14♀, PerMANOVA: P=0.017, F=2.594, R²=0.123). (D) Relative concentrations of CHCs compounds in mated females compared to virgin females. CHCs levels in virgin females were normalized to 1, and the concentrations in mated females were expressed relative to this baseline (LMM and permutation test: n*_virgin_*=10♀, n_Cs-mated_=16♀, n*_Bully-mated_*=14♀, Monte-Carlo test: P=0.618). The absence of a significant multivariate effect does not preclude family-specific or compound-specific differences, as the multivariate tests evaluated the joint profile across all compounds rather than the effect on any family or individual compound. Family-wise comparison (LMM and permutation test P_Alkanes_=0.705, P_Monoenes_=0.294, P_Dienes_=0.715, P_Methyl alkanes_=0.750) and compound-wise comparison (LMM and permutation test: P_nC21_=0.158, Pn_C22_=0.927, P_7,11C23:2_<0.001, P_2MeC22_<0.001, P_C23:1(9)_=0.023, P_C23:1(7)_=0.001, P_C23:1(5)_=0.016, P_nC23_=0.908, P_nC24_=0.899, P_7,11C25:2_<0.001, P_2MeC24_<0.001, P_C25:1(9)_=0.832, P_C25:1(7)_=0.047, P_C25:1(5)_<0.001, P_nC25_=0.769, P_9,13C27:2_<0.001, P_7,11C27:2_=0.747, P_2MeC26_=0.884, P_C27:1(9)_=0.016, P_C27:1(7)_=0.625, P_C27:1(5)_<0.001, P_nC27_=0.367, P_9,13C29:2_<0.001, P_7,11C23:2_=0.899, P_2MeC28_=0.413, P_C29:1(9)_<0.001, P_C29:1(7)_<0.001, P_nC29_=0.022). C23:1(5) and C25:1(5) compounds refer to 5-T and 5-P respectively and are highlighted in the figure as they present a significant difference between virgin, *Cs*- and *Bully*-mated females. Significant differences are indicated by letters: *a* indicates a significant difference between virgin and *Cs*-mated females; *b* indicates a significant difference between virgin and *Bully*-mated females; and *c* indicates a significant difference between *Cs-* and *Bully*-mated females (*a* P < 0.05; *aa* P < 0.01; *aaa* P < 0.001). (E) Mean ± SE concentration in ng of CHCs compounds that were not present in virgin females (n*_virgin_*=10♀, n_Cs-mated_=16♀, n*_Bully-mated_*=14♀, P_C23:1(6)_<0.001, P_C25:1(6)_<0.001, P_cVA_=0.001). (F) Mating duration of *Cs* and *Bully* males with a *Cs* female (n_Cs-mated_=17, n*_Bully-mated_*=14, T-test: P=0.003, t= −3.359). (F’) cVA concentration measured within *Cs*- and *Bully*-mated genitalia tract right after mating (n_Cs-mated_=17, n*_Bully-mated_*=14, Fisher Pitman permutation test: P=0.001, Z= −2.390). For graphs E, F and F’, stars indicate significant differences (* P < 0.05, ** P < 0.01, *** P < 0.001). Post-hoc analyses and statistical details are given in Figure 1-table supplement 1.

**Table 1:**
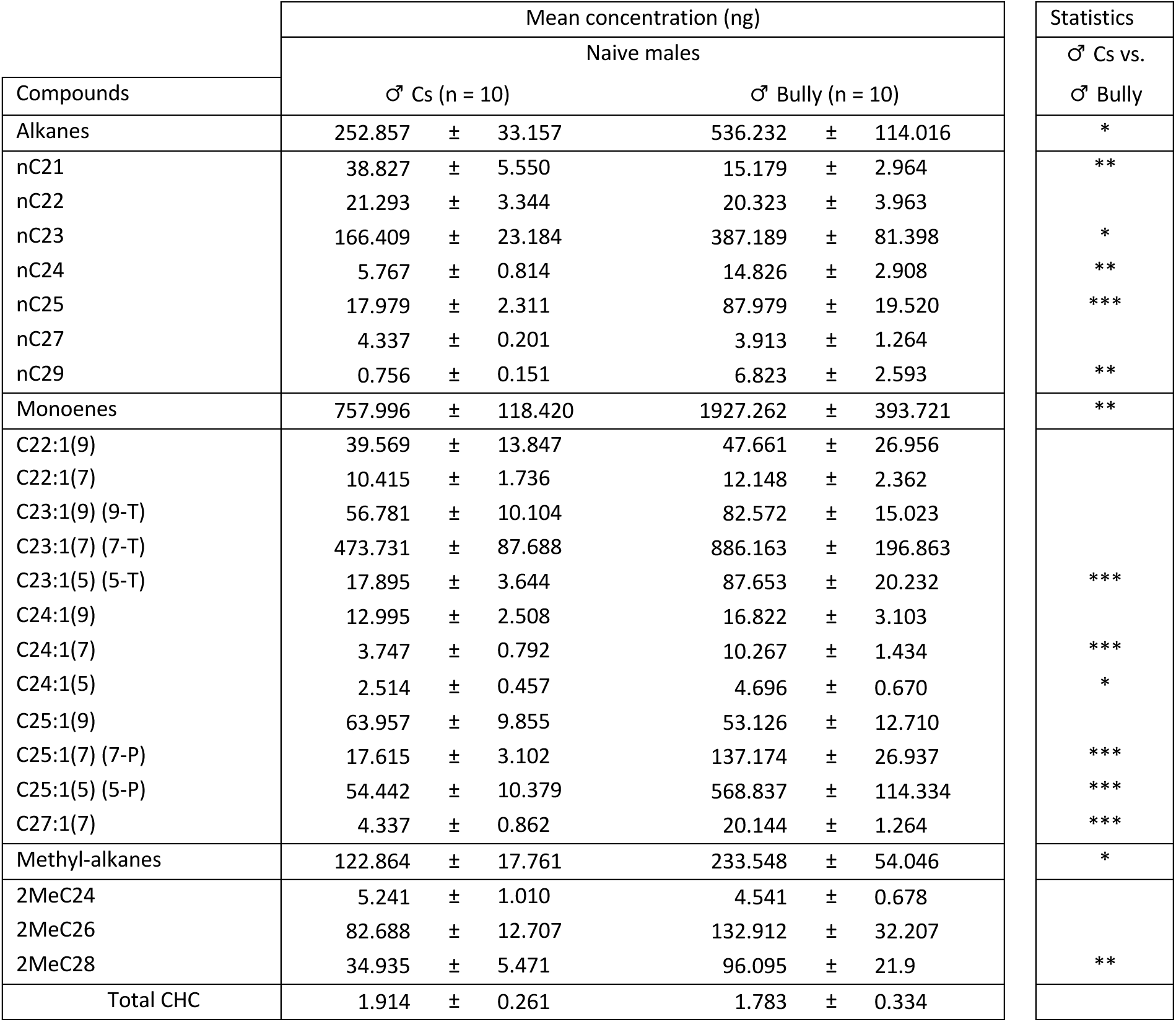
Comparison of. CHCs **concentrations between naïve *Cs* and *Bully* males.** Mean ± SE concentrations are provided in ng for each compound and each group of compounds. In statistics columns, stars represent significant differences (* P < 0.05, ** P < 0.01, *** P < 0.001). Statistical details are given in Figure 1-table supplement 1.

| Compounds | Mean concentration (ng) |  |  |  |  | Statistics |
| --- | --- | --- | --- | --- | --- | --- |
|  | Naive males |  |  |  |  | ♂ Cs vs.<br>♂ Bully |
|  | ♂ Cs (n = 10) |  | ♂ Bully (n = 10) |  |  |  |
| Alkanes | 252.857 | ± 33.157 | 536.232 | ± 114.016 |  | * |
| nC21 | 38.827 | ± 5.550 | 15.179 | ± 2.964 |  | ** |
| nC22 | 21.293 | ± 3.344 | 20.323 | ± 3.963 |  |  |
| nC23 | 166.409 | ± 23.184 | 387.189 | ± 81.398 |  | * |
| nC24 | 5.767 | ± 0.814 | 14.826 | ± 2.908 |  | ** |
| nC25 | 17.979 | ± 2.311 | 87.979 | ± 19.520 |  | *** |
| nC27 | 4.337 | ± 0.201 | 3.913 | ± 1.264 |  |  |
| nC29 | 0.756 | ± 0.151 | 6.823 | ± 2.593 |  | ** |
| Monoenes | 757.996 | ± 118.420 | 1927.262 | ± 393.721 |  | ** |
| C22:1(9) | 39.569 | ± 13.847 | 47.661 | ± 26.956 |  |  |
| C22:1(7) | 10.415 | ± 1.736 | 12.148 | ± 2.362 |  |  |
| C23:1(9) (9-T) | 56.781 | ± 10.104 | 82.572 | ± 15.023 |  |  |
| C23:1(7) (7-T) | 473.731 | ± 87.688 | 886.163 | ± 196.863 |  |  |
| C23:1(5) (5-T) | 17.895 | ± 3.644 | 87.653 | ± 20.232 |  | *** |
| C24:1(9) | 12.995 | ± 2.508 | 16.822 | ± 3.103 |  |  |
| C24:1(7) | 3.747 | ± 0.792 | 10.267 | ± 1.434 |  | *** |
| C24:1(5) | 2.514 | ± 0.457 | 4.696 | ± 0.670 |  | * |
| C25:1(9) | 63.957 | ± 9.855 | 53.126 | ± 12.710 |  |  |
| C25:1(7) (7-P) | 17.615 | ± 3.102 | 137.174 | ± 26.937 |  | *** |
| C25:1(5) (5-P) | 54.442 | ± 10.379 | 568.837 | ± 114.334 |  | *** |
| C27:1(7) | 4.337 | ± 0.862 | 20.144 | ± 1.264 |  | *** |
| Methyl-alkanes | 122.864 | ± 17.761 | 233.548 | ± 54.046 |  | * |
| 2MeC24 | 5.241 | ± 1.010 | 4.541 | ± 0.678 |  |  |
| 2MeC26 | 82.688 | ± 12.707 | 132.912 | ± 32.207 |  |  |
| 2MeC28 | 34.935 | ± 5.471 | 96.095 | ± 21.9 |  | ** |
| Total CHC | 1.914 | ± 0.261 | 1.783 | ± 0.334 |  |  |

We next examined whether mating with *Bully* males induces changes in CHC profiles of females compared to those that have mated with *Cs* males. To test this hypothesis, we extracted CHCs from the cuticles of *Cs* virgin females and *Cs* females mated to either a *Cs* or *Bully* males and compared their CHCs profile. PCA analysis showed that *Cs* virgin females form a distinct cluster from mated females (Figure 4C), indicating that mating induces substantial changes in female CHC profiles. The first two principal components accounted for most of the variance, and several compounds, including C23:1(6), C25:1(6), and cVA, contributed strongly to this separation (Figure 4-figure supplement 1). Consistent with this, individual compound analysis revealed that male-specific CHCs, such as 7-tricosene (7-T), 9-tricosene (9-T), and cVA, were significantly increased in mated females compared to virgins (Figure 4D–E, Table 2). These results are consistent with previous studies showing that 7-T and 9-T are acquired by females during mating (Everaerts et al., 2010; Laturney and Billeter, 2016).

**Table 2:**
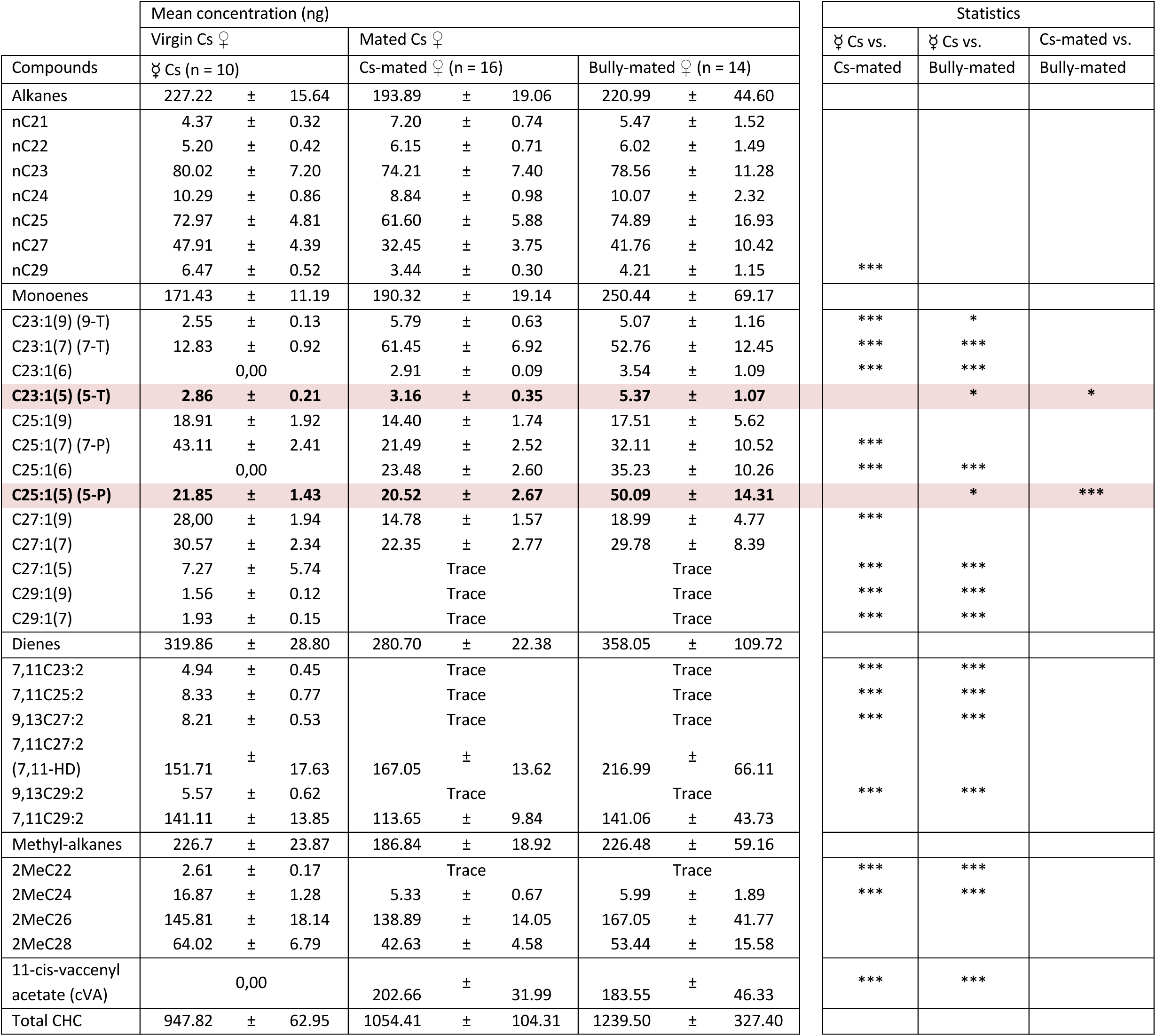
Comparison of. CHCs **concentrations among virgin, *Cs*-mated, and *Bully*-mated females.** Mean ± SE concentrations are provided in ng for each compound and each group of compounds. In the statistics columns, stars indicate significant differences (* P < 0.05, ** P < 0.01, *** P < 0.001). Statistical details are given in Figure 1-table supplement 1.

We then compared CHC profiles between females mated with *Cs* or *Bully* males. While both groups exhibited the expected increase in male-derived compounds, females mated with *Bully* males showed significantly higher levels of 5-tricosene (5-T) and 5-pentacosene (5-P) (Figure 4D, Table 2). As *Bully* males also showed higher levels of both 5-P and 5-T compared to naïve *Cs* males (Figure 4B), it is possible that the elevated levels of these compounds observed in females could result from their transfer during mating.

cVA, a male-specific compound that acts as an anti-aphrodisiac and a mate-guarding molecule, is primarily transferred to the female’s genital tract during mating, with only trace amounts deposited on the cuticle (Laturney and Billeter, 2016). Females with higher cVA levels (Kurtovic et al., 2007), combined with 7-T, are less attractive to males (Laturney and Billeter, 2016). Given that remating rate of *Bully*-mated females was higher than that of *Cs*-mated females (Figure 3G’), we hypothesized that *Bully*-mated females would receive less cVA in their genital tract during mating, thus decreasing their attractiveness than when mating with *Cs* males. To test this, we conducted mating assays with *Cs* females paired with either *Cs* or *Bully* males, followed by dissection of the females’ genital tracts to quantify the amount of cVA transferred. We found that females mated with *Bully* males had about three times lower cVA quantities in their genital tracts compared to those mated with *Cs* males (Figure 4F’). Combined with higher quantities of 5-T and 5-P, reduced cVA level in genital tract were associated with increased attractiveness and higher remating rates observed in females that previously mated with *Bully* males.

Overall, our findings demonstrate that naïve *Bully* males differ from *Cs* males in their CHC profiles and transfer a distinct quantity of chemical compounds to females during mating. These observations are consistent with a model in which changes in chemical signaling in *Bully* males is associated with both reduced mating success and weaker mate-guarding-related behaviors. In particular, reduced cVA transfer, combined with elevated levels of specific hydrocarbons such as 5-T and 5-P in mated females, was associated with increased attractiveness and remating propensity observed in females that previously mated with *Bully* males, although whether these changes are causally related remains unknown.

### Reduced copulatory investment in aggressive males is associated with extended lifespan

Based on the main life-history trade-off between reproduction and survival, we next examined whether reduced pre- and post-copulatory investment observed in *Bully* males would be accompanied by increased longevity. To this end, we conducted longevity assays and monitored lifespan in *Cs* and *Bully A* males. When housed individually, *Bully* males survived significantly longer than *Cs* males (Figure 5A, green vs yellow curves). Median survival (EC50) was increased from 57 days in *Cs* males to 82 days in *Bully* males (Figure 5A), and maximum lifespan was also extended from 90 days in Cs males to 100 days in Bully (Figure 5A). To assess whether this difference could reflect hybrid vigor, we measured lifespan in heterozygous *Bully* males generated by crossing Cs females with *Bully* males. The survival of these heterozygous males was comparable to that of homozygous *Bully* males (Figure 5A, red curve), with males showing an EC50 of 79 days and a maximum lifespan of 106 days. This result indicates that *Bully* background positively influence males’ lifespan and argue against hybrid vigor as the main explanation for the observed phenotype. We next evaluated survival under more naturalistic social conditions by maintaining males in mixed-sex groups. Under these conditions, *Bully* males again showed increased survival compared to *Cs* males, with both higher median lifespan and extended maximum lifespan (Figure 5B). *Bully* males had a median survival of 56 days and a maximum lifespan of 71 days, compared to *Cs* males, which had a median survival of 43 days and all died by day 67 (Figure 5B).

**Figure 5.**
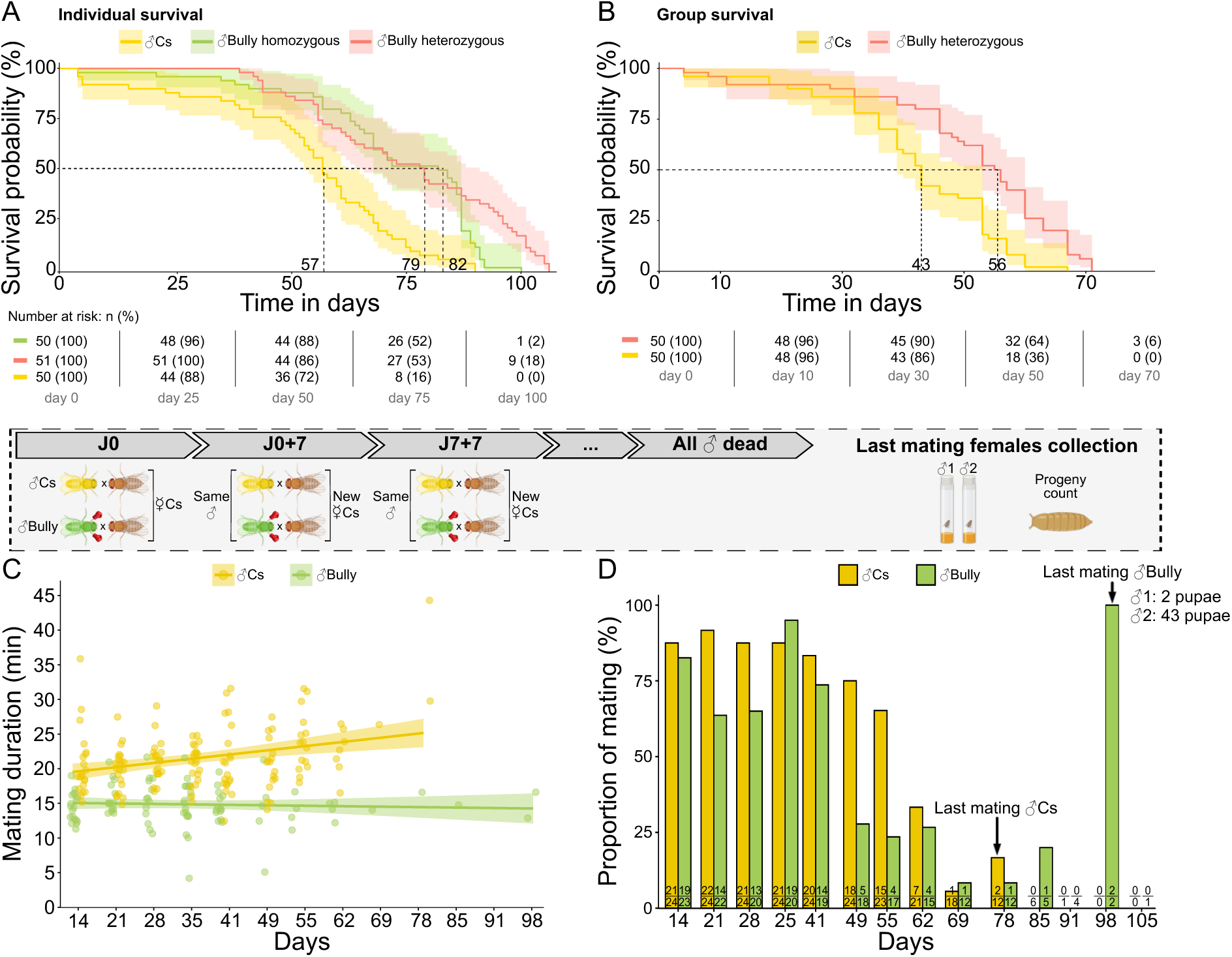
Aggressive males offset their reduced reproductive success with a longer lifespan and preserved reproductive success in later life. (A) Survival curves of *Cs* (yellow) and homozygous (green) and heterozygous (red) *Bully* males (Weibull regression model (n_Cs_=50, n*_Bully-homo_*=50, n_Bully-hetero_=51, Likelihood Ratio Test): χ²=23.91, p=2.6×10^−5^). *Cs* males had a median survival (EC₅₀) of 57 days, with the last individual dying at day 90 (yellow), while progeny from homozygous *Bully* males showed an EC₅₀ of 82 days, with the final death at day 101 (green) and heterozygous *Bully* males showed an EC₅₀ of 79 days, with the final death at day 106 (red). (B) Survival curves of group of 10 *Cs* and 10 homozygous *Bully* males raised in group with 12 *Cs* females. *Bully* males had a median survival (EC₅₀) of 56 days, with the last individual dying at day 71 (red line), while *Cs* males showed an EC₅₀ of 43 days, with the final death at day 67 (yellow line) (n_Cs_=50, n_Bully-hetero_=50, Cox proportional hazards regression model with stratification by group (Likelihood Ratio Test): P=7×10^−5^, χ²=15.94). (C) Mating duration of *Cs* and *Bully* males with virgin *Cs* females across their lifespan (n_Cs-J0_=24, n_Bully-J14_=23, LMM and Satterthwaite T-test: P_interaction_=3.360×10^−5^, t=4.227). A significant interaction between male genotype and age was observed: *Bully* males maintained a stable mating duration over time, whereas *Cs* males showed an increase in mating duration with successive mating events. (D) Number of mating throughout the lifespan of *Cs* and *Bully* males (n_Cs-J0_=24, n_Bully-J14_=23, GLM binomial and Wald Z-test: P_interaction_=0.039, Z= −2.068). Numbers in bars indicate the proportion of mating. Arrows denote the last mating event and the number of offspring produced. Post-hoc analyses and statistical details are given in Figure 1-table supplement 1.

We then assessed how reproductive performance varies across the lifespan. Individual males were paired weekly with virgin *Cs* females until death, and mating success and mating duration were recorded. Throughout their lifespan, *Bully A* males consistently displayed shorter MD than *Cs* males (Figure 5C). While mating duration increased with age in Cs males, a pattern reported in another *Drosophila* species (Dhole and Pfennig, 2014), this effect was not observed in *Bully* males, whose MD remained stable across mating and throughout their lives (Figure 5C, green). Analysis of individual male slopes confirmed this divergence: most *Cs* males showed increasing MD across mating events, while *Bully* males did not (Figure 5-figure supplement 1). Mating success followed a similar age-dependent pattern. *Bully* males showed reduced mating success compared to *Cs* males during early and mid-adulthood (until 70 days), but continued to mate at ages when *Cs* males had ceased reproductive activity (Figure 5D). While all *Cs* males stopped mating by day 78 and died by day 97 (Figure 5D, yellow), *Bully* males maintained mating activity until day 98 and survived up to 119 days (Figure 5D, green). In addition, late-life mating involving 98-days-old *Bully* males produced viable offspring (Figure 5D, green), demonstrating that *Bully* males remain fertile at older ages.

Together, our findings demonstrate that *Bully* males exhibit both reduced copulatory investment and extended lifespan, as well as a shift in the temporal pattern of reproductive activity. These observations are consistent with a scenario in which reduced investment in mating success and mating duration is associated with extended lifespan and sustained reproductive output later in life. Rather than enhancing early-life reproductive success, *Bully* males maintain reproductive capacity over a longer time window, suggesting a redistribution of reproductive effort across the lifespan.

## DISCUSSION

Our study reveals that selection for aggressive traits in *D. mel.* males correlates with changes in key life-history trade-offs, in which survival is extended while reproductive success is reduced. This pattern challenges the hypothesis that aggression necessarily enhances mating success (Andersson, 1994; West-Eberhard, 1983), and indicates that aggressive phenotypes may incur substantial reproductive costs. While the behavioral phenotypes were replicated across three independently selected Bully lines, the evidence for the survival phenotype comes from *Bully A* males only. In this line, increased lifespan was associated with reduced reproductive success and a shift in the timing of reproductive activity. Testing lifespan in additional *Bully* lines will be necessary to determine whether this association between aggression selection and lifespan is shared across independently selected lines or reflects line-specific changes. Behavioral phenotypes were replicated across distinct *Bully* lines, although *Bully C* males showed an intermediate increase of aggression and no reduction in mating duration. This pattern could reflect graded strength of the selected phenotype or line-specific changes, including attenuation of the initial aggressive phenotype in *Bully C*. Further experiments would be required to disentangle between these possibilities.

The absence of additional selected lines limits our ability to fully exclude contributions of genetic drift or line-specific effects. Furthermore, because the *Bully* and *Cs* lines have been maintained independently for many years following the original selection experiment, we cannot exclude that additional genetic divergence accumulated during laboratory maintenance contributed to some of the observed phenotypes. Because selection for aggression was applied specifically to males, we used a standardized female background (*Cs*) to reduce variability arising from female genotype. While this approach facilitates comparison of male phenotypes, it does not fully eliminate potential interactions between male and female genotypes. The persistence of mating duration phenotype in *Bully* × *Bully* pairings and across independent selected lines, supports a primary contribution of male genotype, but does not establish whether female genotype also contributes to this phenotype or interacts with male genotype. Reciprocal crosses and factorial combinations of male and female genotypes would be required to disentangle these effects, particularly for reproductive and chemical phenotypes and for lifespan under social or mating conditions.

Aggression therefore does not act as an isolated fitness-enhancing trait, but rather as one component of a coordinated life-history strategy in which gains in competitive ability are coupled to constraints on reproductive performance. The increased survival of highly aggressive males further raises important questions regarding the evolutionary consequences of aggression. While *Bully A* males have reduced mating success compared to *Cs* males, their extended lifespan may allow them to sustain mating activity over longer periods and produce viable offspring at later ages, challenging the classical view that early reproductive success is the primary driver of male fitness (Flatt, 2011). Reduced mating frequency could contribute to the survival difference under group-housed conditions, although the longer lifespan of isolated *Bully A* males, which never mate, indicates that reduced mating is not sufficient to explain the increased longevity. This shift suggests that alternative reproductive strategies, even those associated with lower mating success and mate-guarding, can persist if they confer long-term survival advantages under laboratory conditions.

At the behavioral level, highly aggressive *Bully A* males exhibit changes in both pre- and post-copulatory phases. They achieve lower mating success and engage in shorter copulations than less aggressive *Cs* males, suggesting reduced efficiency in mate acquisition and potentially in sperm transfer. Despite higher courtship motivation after mating, *Bully* males do not achieve higher remating success. Instead, females previously mated with these males remate more readily, indicating reduced mate-guarding efficiency and possibly increased attractiveness. These observations point to a decoupling between courtship intensity and mating outcome, indicating that increased behavioral drive does not translate into reproductive advantage. While our study did not show a causal link, our results suggest that artificial selection applied to male aggressive traits to produce the *Bully* A line, is associated with changes in reproductive traits, enhanced survival, and increased CHCs levels.

These behavioral differences are closely associated with significant modifications in chemical signaling. Cuticular hydrocarbons (CHCs), which are crucial to mate recognition and sexual communication (Billeter and Wolfner, 2018; Yew and Chung, 2017) differ significantly between *Bully A* and *Cs* males. We show that naïve highly aggressive individuals present distinct chemical profiles compared to unselected wild-type flies, with higher levels of most CHCs. Importantly, CHCs also contribute to physiological functions such as protection against desiccation (Nayal et al., 2024), suggesting that changes in CHC composition may simultaneously influence both reproductive performances and survival. Thus, selection on aggressive traits may have been accompanied by modifications in overall chemical signaling that would impact both physiological resilience and social behaviors.

Although higher CHC level might be expected to enhance male attractiveness, *Bully* males display reduced mating success, suggesting a non-linear or dose-dependent relationship between CHC levels and female acceptance. Female mate choice relies on finely tuned blends of chemical cues, and changes in these optimal ratios may impair signal interpretation (Billeter and Wolfner, 2018). In this context, excessive levels of specific CHCs could reduce courtship efficiency rather than enhance it. In addition to major compounds such as 7-tricosene, male attractiveness and courtship performance are influenced by other CHC, including palmitoleic acid, which signals through the Or47b receptor and modulates courtship behavior in an age-dependent manner (Dweck et al., 2015; Kohlmeier et al., 2021; Lin et al., 2016). This pathway enables older males to initiate courtship more rapidly and with greater intensity, potentially compensating for age-related declines in competitiveness. Although palmitoleic acid levels were not measured here, change in level of such compounds could further contribute to decreasing mating success despite overall higher CHC level.

Modifications in chemical communication extend into the post-mating context, where they potentially intersect with male reproductive performance. Females mated with *Bully* males display distinct CHC profiles, with higher levels of 5-tricosene (5-T) and 5-pentacosene (5-P), together with reduced transfer of cis-vaccenyl acetate (cVA) into the female genital tract, which may fail to efficiently decrease their attractiveness, thus triggering higher remating rate. This interpretation is consistent with previous studies showing that lower cVA levels decrease the effectiveness of chemical mate guarding and accelerate the recovery of female attractiveness (Ejima, 2015). At the same time, the behavioral effects of cVA depend on its interaction with other CHCs (Laturney and Billeter, 2016; Verschut et al., 2023), suggesting that changes in the overall chemical profile may further modulate how mated females are perceived by subsequent males. Thus, changes in chemical signaling is a candidate mechanism to link aggression selection to post-copulatory reproductive traits, but a causal relationship remains to be established. CHC and behavioral changes could also represent independent consequences of genetic changes associated with selection for aggression. Direct manipulation of CHCs or cVA will be required to distinguish between these possibilities.

Taken together, these observations support the idea that cuticular hydrocarbons constitute a central mechanistic substrate linking aggression, reproduction, and survival. Rather than hypothezing selection on multiple behavioral traits, the phenotypic differences observed in our study may alternatively arise from selection acting on genes involved in chemical signaling. Because CHCs influence both social communication and physiological resilience (Nayal et al., 2024), selection that has acted on CHC production or emission would be expected to generate correlated effects across social behavior, reproductive success, and survival, and physiological resilience, thereby producing the unique life-history pattern observed in *Bully* males.

At the genetic level, this integration may be supported by pleiotropic factors or by the co-selection of partially independent loci affecting multiple traits. In this study, we mainly focused on *Bully A* males, as transcriptomic analyses have identified a limited set of genes differentially expressed between *Bully A* and *Cs* males, including *CG13646*, a putative transmembrane transporter whose down-regulation increases aggressive behavior (Chowdhury et al., 2017). These candidate genes provide an entry point for dissecting the molecular and genetic logic underlying the observed phenotypes, particularly in determining whether changes in aggression, CHC composition, survival and life-history traits arise from common genetic regulation.

In summary, our findings indicate that selection for male aggression is associated with a coordinated shift in reproductive investment, chemical signaling and lifespan, consistent with a redistribution of reproductive effort across the lifespan. This highlights aggression as a potential entry point into the coordinated evolution of reproductive and physiological traits.

### MATERIAL AND METHODS

### EXPERIMENTAL MODEL AND SUBJECT DETAILS

#### Fly stocks

*Drosophila melanogaster* strains were raised at 25°C under a 12h:12h light/dark cycle (LD= 9:00a.m. – 9:00p.m.) on standard fly food (for 1l of water, 74g corn flour, 28g yeast, 40g sugar, 7.5g agar, 10ml Moldex). Canton-S (*Cs*) flies (from Edward Kravitz’s laboratory, RRID: BDSC_64349) were used as wild-type. The *Bully* A, B and C lines were originally selected in Edward Kravitz’s laboratory from *Cs* during 37 generations (Penn et al., 2010), resulting in hyper-aggressive male flies (Garbaczewska et al., 2013; Hopkins et al., 2019).

#### Experimental arena

Experimental arenas were described in details in Trannoy *et al*., 2015 (Trannoy et al., 2015a). Briefly, resin blocks containing three circular behavioral arenas (dimension: 2.3 cm diameter, 1.7 cm height) were used to conduct behavioral assays. Each arena was divided in two equal sizes by a plastic divider allowing flies to acclimate without physical interactions with the second fly. Then, dividers are removed from the arenas to start the behavioral assays. All behavioral assays were performed in empty arenas, except for aggression assays and courtship assays with decapitated females, in which a food cup (1.5 cm diameter, 1 cm height) containing fresh standard fly food with a drop of fresh yeast paste on the surface was placed in the center of each arena.

### METHOD DETAILS

#### Commonalities between all behavioral assays

Late-stage pupae were collected from stocks raised at 25°C. Male pupae were individually placed in 5ml vials (Dutscher, ref 390597) containing about 1ml of standard fly food while female pupae were placed in group of 15 in ≈47ml vials (Dutscher, ref 789001B) containing about 15ml of standard fly food. All vials were placed at 25°C for 7 days. Seven-day-old individuals were randomly inserted in arenas by negative geotaxis, and allowed to acclimate for 5 min before experiments. All behavioral assays were performed at 25°C. All tests were recorded with Basler camera Ace1970 through the StreamPix8 multi video recording software (norpix.com) and analyzed with BORIS software (Friard and Gamba, 2016). After 15 min or 30 min, if pairs of flies had not yet lunged or mated respectively, we considered that no aggression or mating occurred and arenas were not considered for analysis. No power analyses were performed prior to the experiments, as we chose to use sample sizes of approximately 20 replicates, which is considered standard practice in the field.

#### Aggression assays

Two days prior to the aggression assays, males were anesthetized with CO₂, and half were marked with a small white dot on the thorax before being returned to their original isolation vials. Aggression tests were conducted between ZT0 and ZT3 using size-matched male pairs. Males competed for access to a central food cup containing yeast. The following behaviors were recorded for 10 minutes following the first lunge: latency to lunge (i.e., the time between the initial encounter on the food cup and the first lunge), total number of lunges, and number of boxing events.

#### Mating assays

Seven-day-old individuals were tested in arenas without any food source. Latency to court and to mate, defined as the time between the first encounter and either the first Unilateral Wing Extension (UWE) or the initiation of mating, respectively, were measured. Total UWE duration was recorded over a 10-minute period starting from the first encounter. Mating proportion (proportion of fly pairs that succeeded to mate) and mating duration were also scored. In all assays, males were reared in social isolation, except for Figure 1H, where they were housed in groups of 10 males for 7 days.

#### Mating assays with decapitated females

Females were anesthetized with CO₂ and placed under the binocular for head removal. Experiments with decapitated females were performed in arenas with food cup filled with fresh fly food but without yeast. The females’ bodies were gently pressed into the food cup to ensure they remained completely immobile, as males can move their bodies during courtship. This allows to maintain a constant distance between the two decapitated females across trials, thereby improving the reproducibility of the assay. Female bodies were positioned in the food cup with the abdomen raised, allowing males to initiate and complete copulation.

#### Multiple consecutive mating events assays

Single naïve *Cs* and *Bully* males were sequentially paired with six virgin *Cs* females. For each male, number of mating and each mating duration were recorded. After each mating, the mated female was transferred to a 5 ml vial containing fly food to assess progeny production.

#### Mating assays throughout the lifespan

24 *Cs* and *Bully* males were reared in isolation and paired with seven-day-old virgin *Cs* females for mating assays, conducted every seven days until all males had died. At each time point, mating duration and mating success (proportion of mating) were recorded. After all *Cs* males had died, females that had mated with *Bully* males were transferred to 5 ml vials containing fly food to assess progeny production.

#### Attractiveness assays

#### Measure of post-mating UWE

Mating assays were performed by pairing naïve *Cs* and *Bully* males with a *Cs* female. Right after mating end, total time that males displayed UWE for 10 minutes towards the just-mated females was scored.

#### Measure of post-mating UWE with switched females

Mating assays were performed with naive *Cs* or *Bully* males and virgin *Cs* females. Immediately after mating ends, females were switched: *Cs* males interacted with *Bully*-mated females and *Bully* males interacted with *Cs*-mated females. Then total time of UWE towards the mated females was scored during 10 min.

#### Measure of post-mating UWE with decapitated females

Mating assays were conducted using naïve *Cs* or *Bully* males paired with virgin *Cs* females. Following copulation, mated females were decapitated. One decapitated *Cs*-mated and one *Bully*-mated female were then placed on the surface of the food cup. A new naïve *Cs* or *Bully* male was introduced and allowed to interact with both decapitated females. The duration of Unilateral Wing Extensions (UWE) directed toward each female was recorded over a 2-minute period following the first encounter. Arenas in which males interacted with only one of the two females were excluded from the analysis, as no choice could be determined.

#### Males’ perception assays

A decapitated *Cs* male and female were placed on a food cup in presence of either a naïve *Cs* or a *Bully* male. The time spent by the focal male courting each decapitated fly was quantified over a 2-minute period following the first unilateral wing extension (UWE). Arenas in which the male interacted with only one of the two decapitated flies were excluded from the analysis, as no choice could be determined.

#### Re-mating assays

Mating assays were performed with naïve *Cs* or *Bully* males and virgin *Cs* females. Mated-females were then individually placed in 5ml vials containing about 1ml of standard fly food and kept at 25°C. 12 or 14 days after the initial mating, mated-females were paired with a new naïve *Cs* male in empty arena and the proportion of re-mating during 30 min was scored.

#### Survival assays

#### Survival with isolated males

*Cs*, homozygous and heterozygous *Bully* late-stage pupae were individually isolated in 5 ml vials containing 1ml of standard fly food and kept at 25 °C until adult emergence. The number of dead individuals was recorded daily until all flies had died. To maintain food quality, flies were transferred weekly to fresh vials containing approximately 1 ml of new fly food.

#### Survival with group-raised males

Late-stage *Cs and* homozygous *Bully* male pupae were placed in group with *Cs* female pupae in ≈47ml vials containing 15mL standard fly food and kept at 25 °C until adult emergence. Groups were composed of 10 males (either *Cs* or *Bully*) and 12 *Cs* females. The number of dead males was recorded daily until they all had died. To maintain food quality, flies were transferred twice a week to fresh vials containing approximately 15 mL of new fly food.

#### CHCs analysis

Extraction and analysis of cuticular hydrocarbons (CHCs) of naive *Cs* and *Bully* males, as well as virgin *Cs*, *Cs*- and *Bully*-mated females were performed in Jean-Christophe Billeter’s lab, Groningen, Netherlands, with a Gas Chromatography coupled with Flame Ionization Detection (GC-FID) as already described (Laturney and Billeter, 2016). Briefly, single whole flies were put in a 2ml glass vial (Supelco certified vial kit, 2ml clear glass vial 12×32mm and Supelco inserts) containing 50µl of nC18+nC26 10 ng/µl standard solution (made in Billeter’s lab) diluted in n-Hexane 99%+ (Acrōs Organic®). Vials were then vortexed at the lowest speed for 2min, afterward flies’ body were gently removed from vials using a “J-shape” paper clip sliced at the end, taking care not to damage the body to prevent from hemolymph contamination. Finally, samples were placed in GC-FID for apolar compounds analysis only.

#### cVA quantification into genital tract

Mating assays were performed with naïve *Cs* or *Bully* males and virgin *Cs* female. Right after mating, females were transferred into a 5ml vial previously put in ice in −80°C fridge to make them fall asleep quickly and prevent from sperm ejection. Mated-females’ genital tract was dissected in PBS with precision forceps under a binocular loupe. Protocol was already employed in (Verschut et al., 2022). Briefly, abdomen was torn to make ovaries and genital tract out and visible, then ovaries, oviduct, spermathecae, parovaria and cuticula were removed to keep only the uterus, taking care not to injure it to prevent from sperm leak. Each uterus was inserted in a 2ml glass vial (Supelco certified vial kit, 2ml clear glass vial 12×32mm and Supelco inserts) containing 50µl of n-Hexane 99%+ (Acrōs Organic) for 1h at ambient temperature. Hexane solution was then evaporated under helium flow to prevent from cVA oxidation with dioxygen. Samples were sent to Jean-Christophe Billeter’s lab for GC-FID analysis of cVA, as describe above in CHCs analysis method part.

### QUANTIFICATION AND STATISTICAL ANALYSIS

No data have been removed from the statistical analysis (except arenas where no choice was observed within the males’ perception assays, as described below). Some experiments were independently replicated by different authors to reduce potential bias. All statistical analyses were performed on R software version 4.0. (Team, 2020) with packages (Fox and Weisberg, 2019), coin (Hothorn et al., 2006), dplyr (Wickham et al., 2023), factoextra (Kassambara and Mundt, 2020), FactoMineR (Lê et al., 2008), lme4 (Bates et al., 2015), lmerTest (Kuznetsova et al., 2017), MASS (Venables and Ripley, 2002), MuMIn (Bartoń, 2023), nlme (Pinheiro et al., 2022), performance (Lüdecke et al., 2021), pgirmess (Giraudoux. P, 2018), predictmeans (Luo et al., 2021), survival (Therneau, 2022), vegan (Oksanen J, 2022).

Comparisons of means (k =2) were performed with a T-test when possible (*i.e.* mating duration) otherwise a Fisher Pitman permutation test was used (*i.e.* UWE, latency to court and to mate). For comparisons with more than 2 conditions, a linear model (LM) was used and tested either with a type II ANOVA when possible (*i.e.* mating duration) or a permutation test instead (*i.e.* latency to lunge and CHC concentrations). For comparison of proportions and counting events, a type II analysis of deviance table with a Likelihood Ratio Test (*i.e.* mating success, proportion of female re-mating, number of lunge and boxing events) or a Wald Chi-square test (*i.e.* number of consecutive mating events) was performed. A Principal Component Analysis (PCA) was performed to see different CHC profiles and tested with a PerMANOVA and a SIMPER analysis. Survival data were analysed with a Weibull regression model and a Likelihood Ratio Test.

When paired data, dependency was considered thanks to linear mixed model (LMM) with the source of dependency as random factor. Data following a distribution other than a normal distribution (*e.g.* binomial or Poisson distribution) were computed in a generalized linear model (GLM) (*i.e.* mating success, number of consecutive mating events, number of mating giving progeny, proportion of females re-mating, number of lunges and boxing events). When necessary, interactions were considered within models and tested. All post-hoc comparisons were performed with a sequential Bonferroni correction on the alpha level using either Holm method (Holm, 1979) when few numbers of comparisons or Benjamini and Hochberg method (Benjamini and Hochberg, 1995) when more comparisons. All statistical details are given in Figure 1-table supplement 1.

## ACKNOWLEDGEMENTS

We thank the members of EXPLAIN and IVEP team at the CRCA for their helpful discussion and support on this study. We thank Kravitz’s lab for sharing the *Bully* lines. We also thank all the member of Billeter’s lab to host A. Defert and train him to GC-FID methodology. We acknowledge support from the French Research National Agency (ANR) (ANR-19-CE37-0018-01 to S.T.), the Fondation Fyssen (190573 to S.T.), and the Doctoral Mobility Campaign of Toulouse III University in 2023 (to A.D.). Funders had no role in study design, data collection and analysis, decision to publish or preparation of the manuscript.

## RESOURCE AVAILABILITY

Further information and requests for resources and reagents are accessible from the lead contact, Séverine Trannoy.

## AUTHOR CONTRIBUTIONS

Conceptualization: S.T., J.C.B., A.D.

Data curation: A.D., R.G., G.P., F.J., A.C., A.H., T.G., S.T.

Formal analysis: A.D., R.G., G.P., F.J., A.C., A.H., T.G., S.T.

Funding acquisition: S.T., J.C.B., A.D.

Project administration: S.T.

Supervision: S.T., J.C.B.

Writing: S.T., J.C.B., A.D.

## SUPPORTING INFORMATIONS

**Figure 1-table supplement 1.**
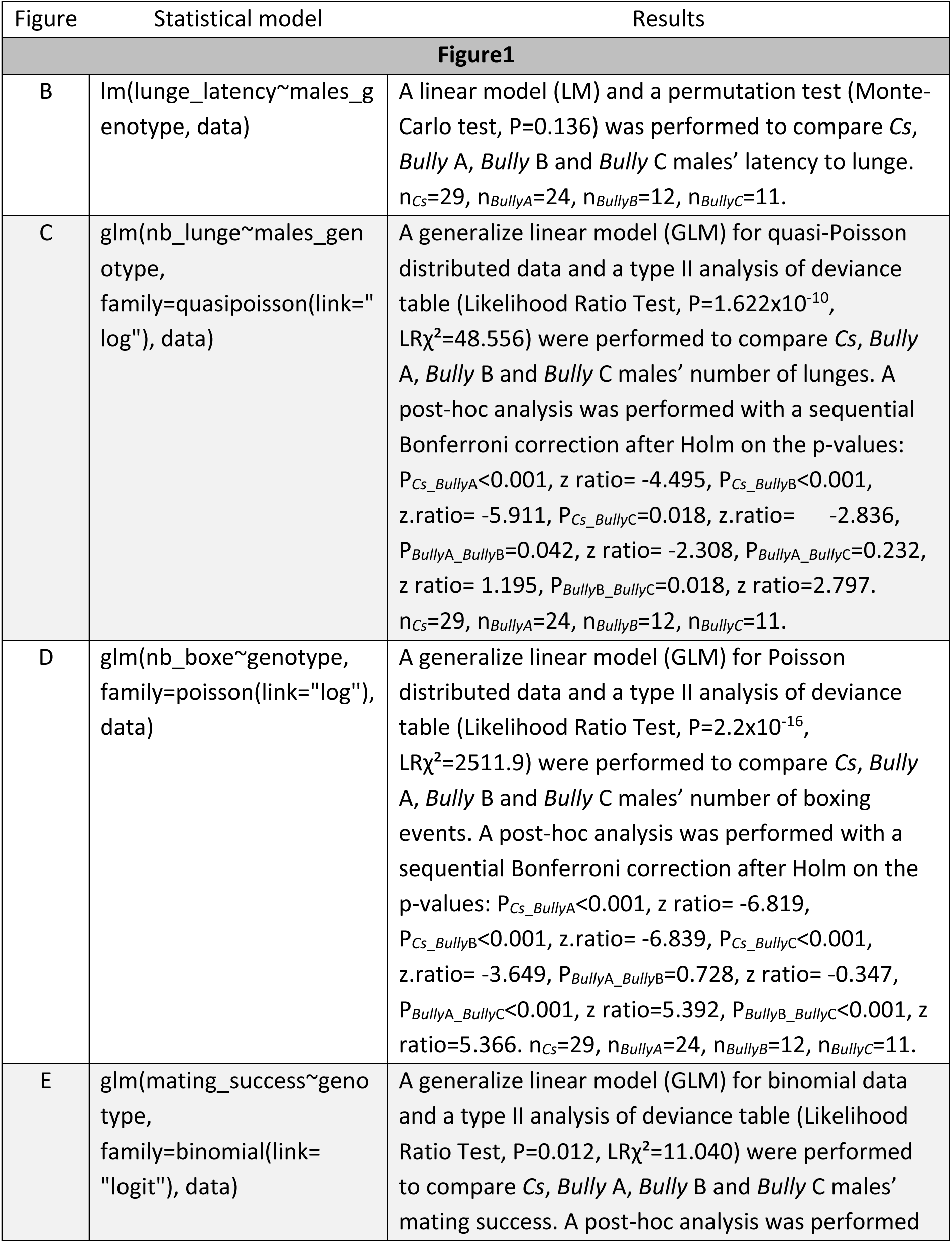
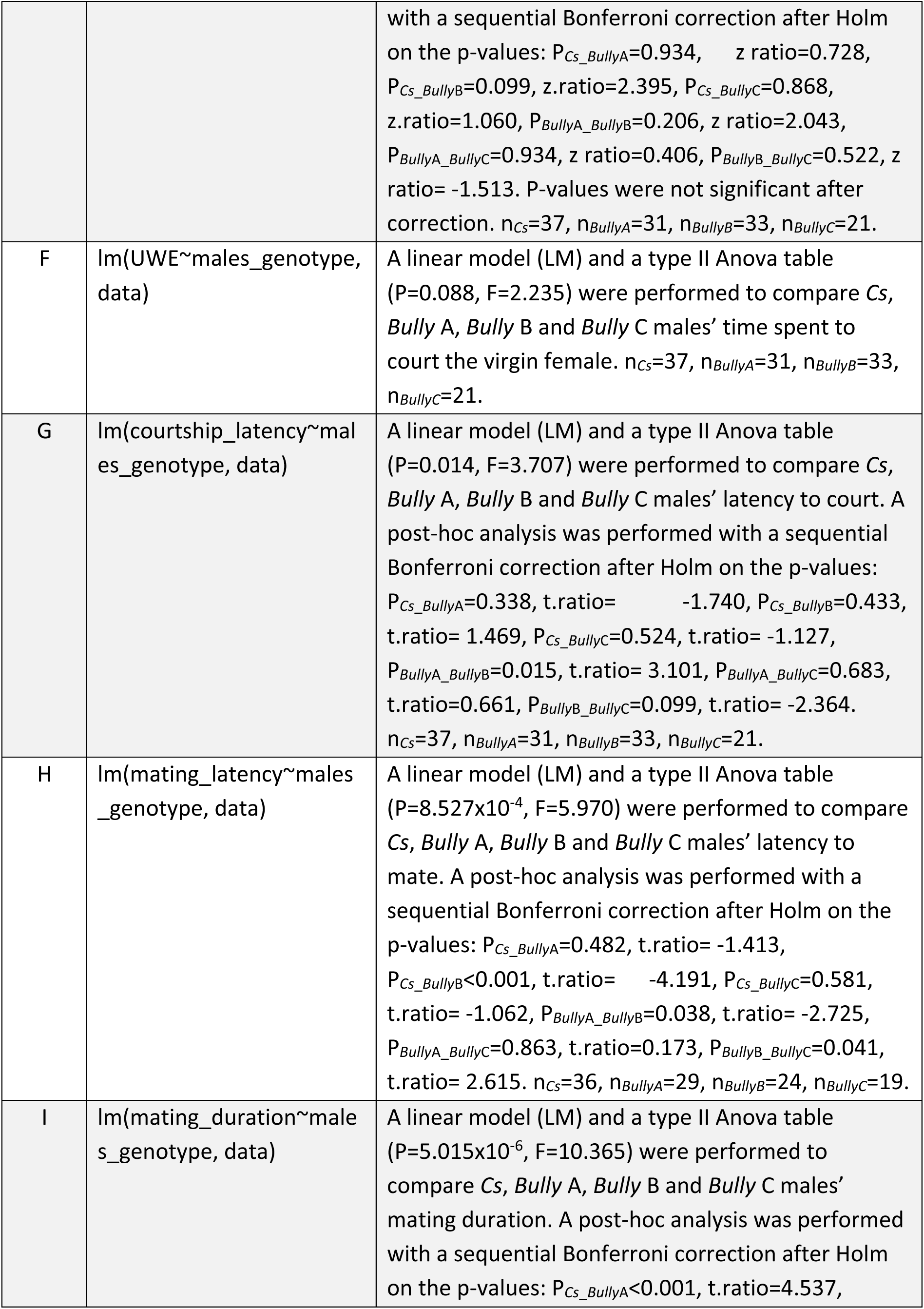
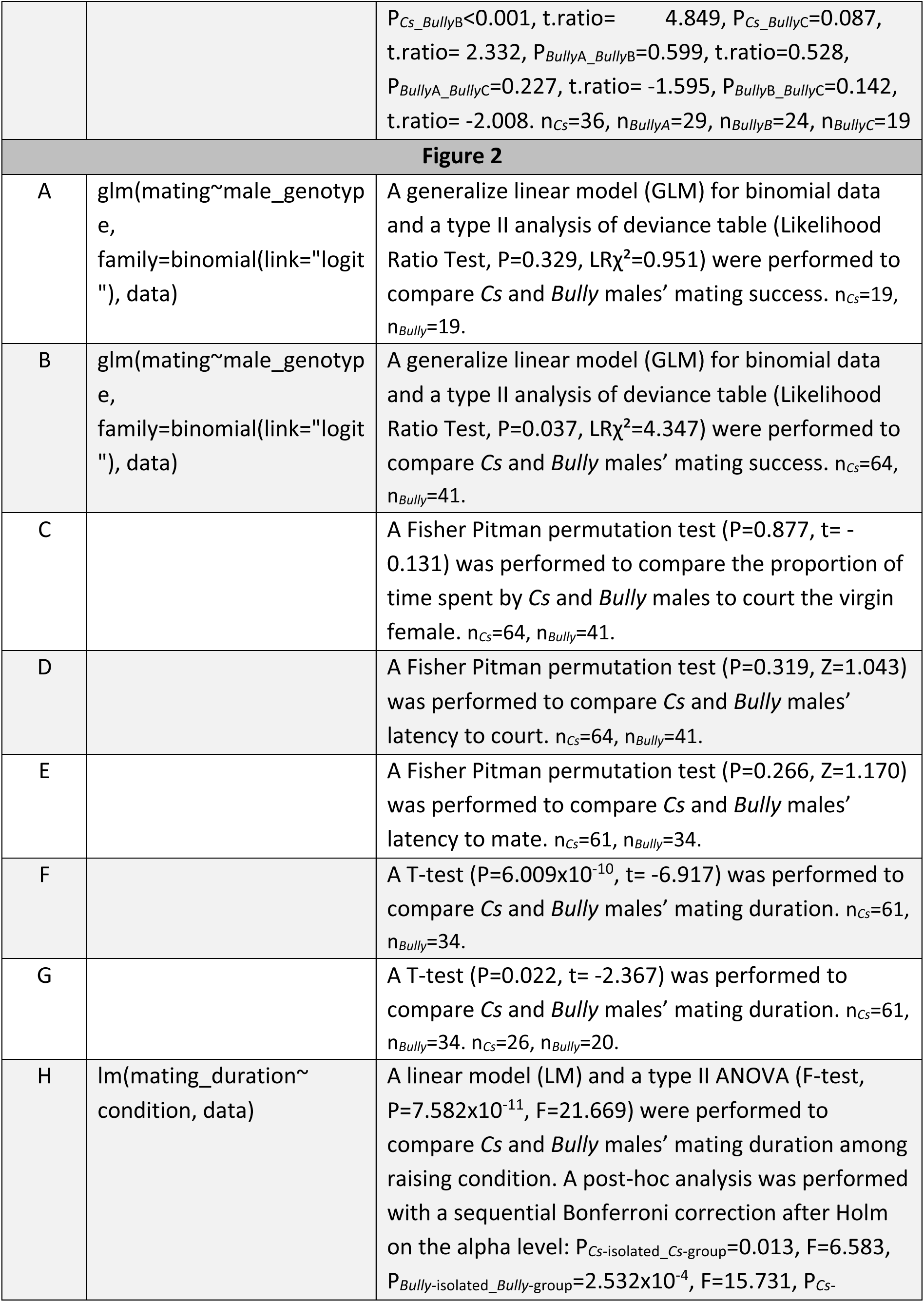
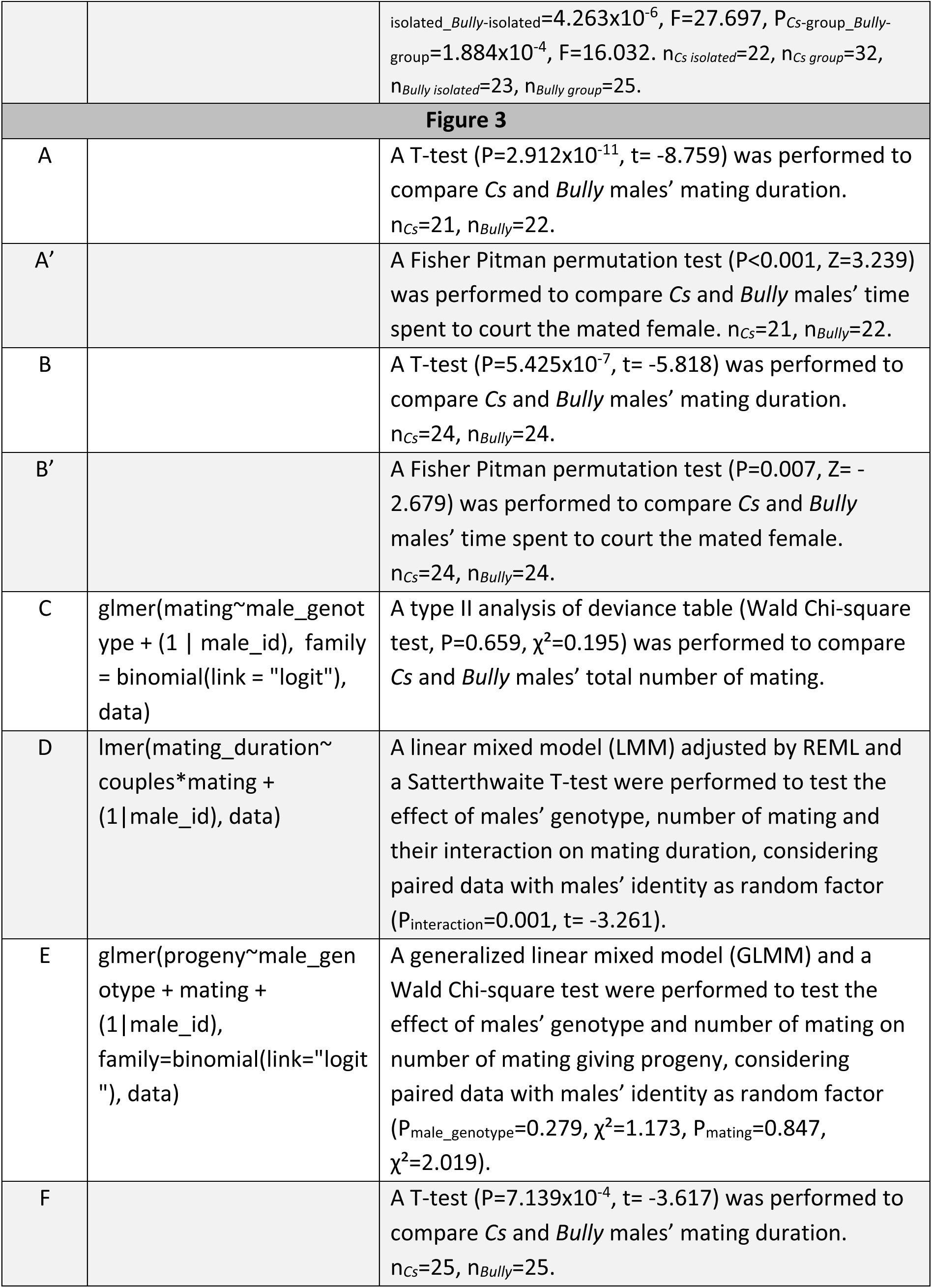
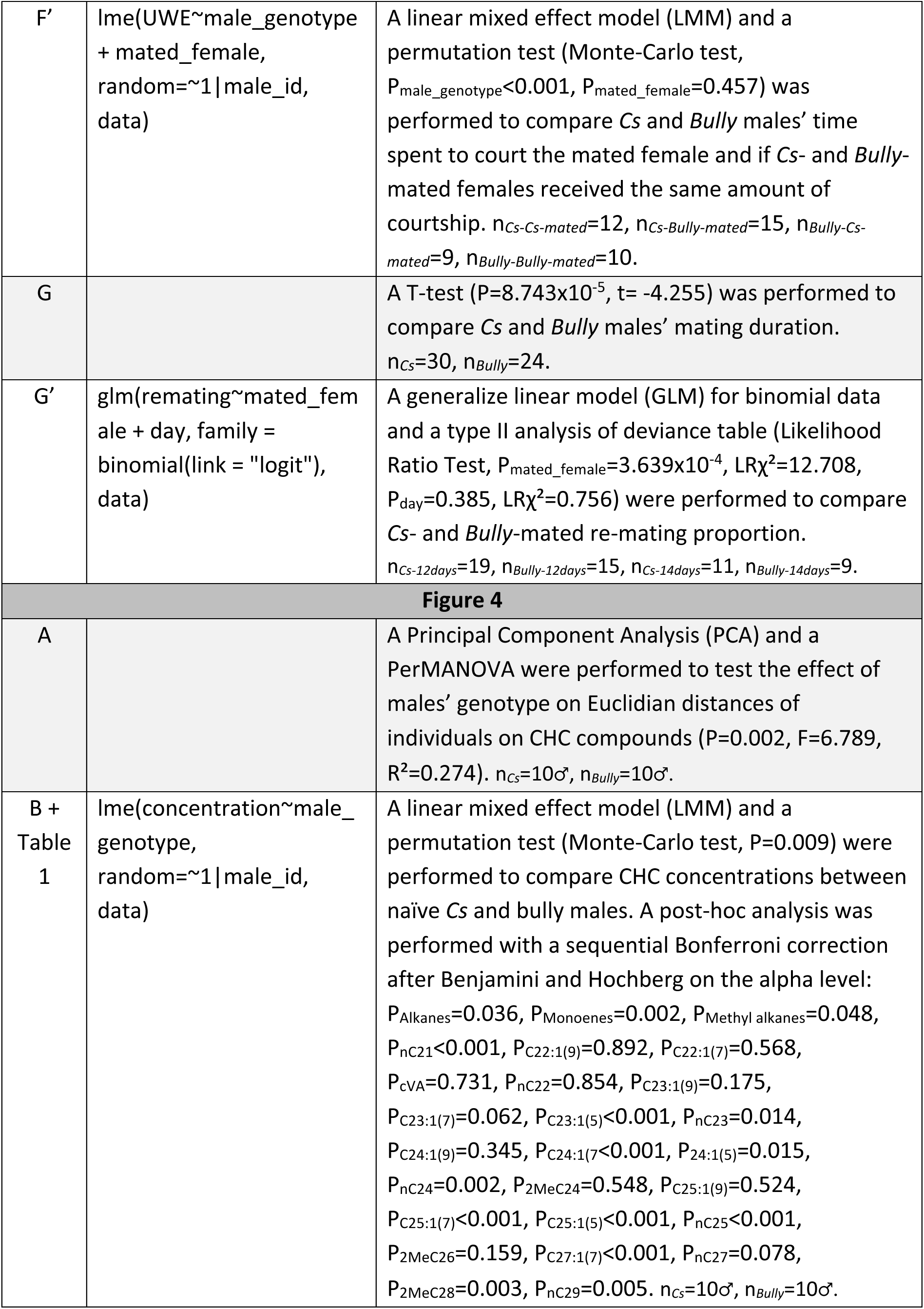
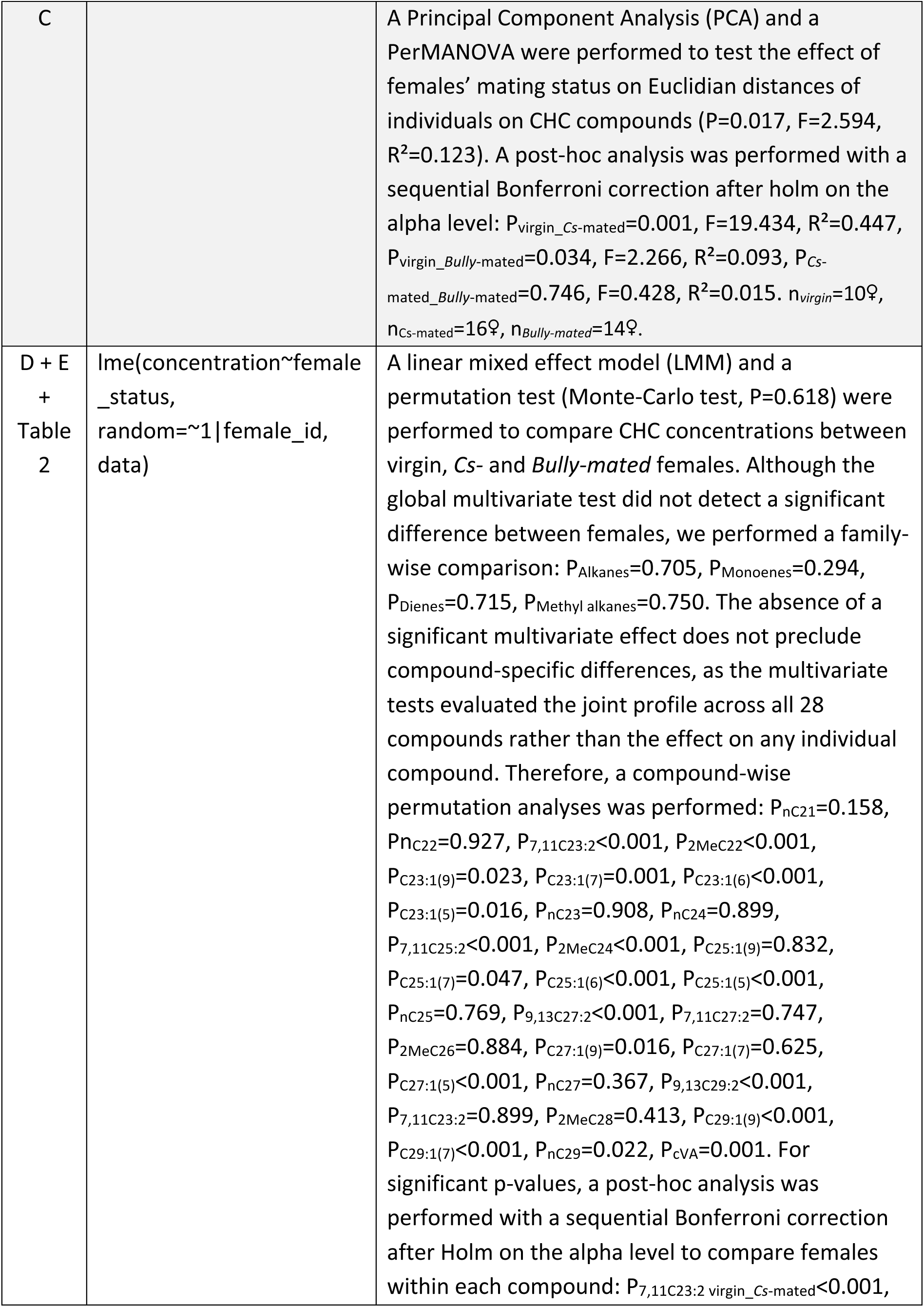
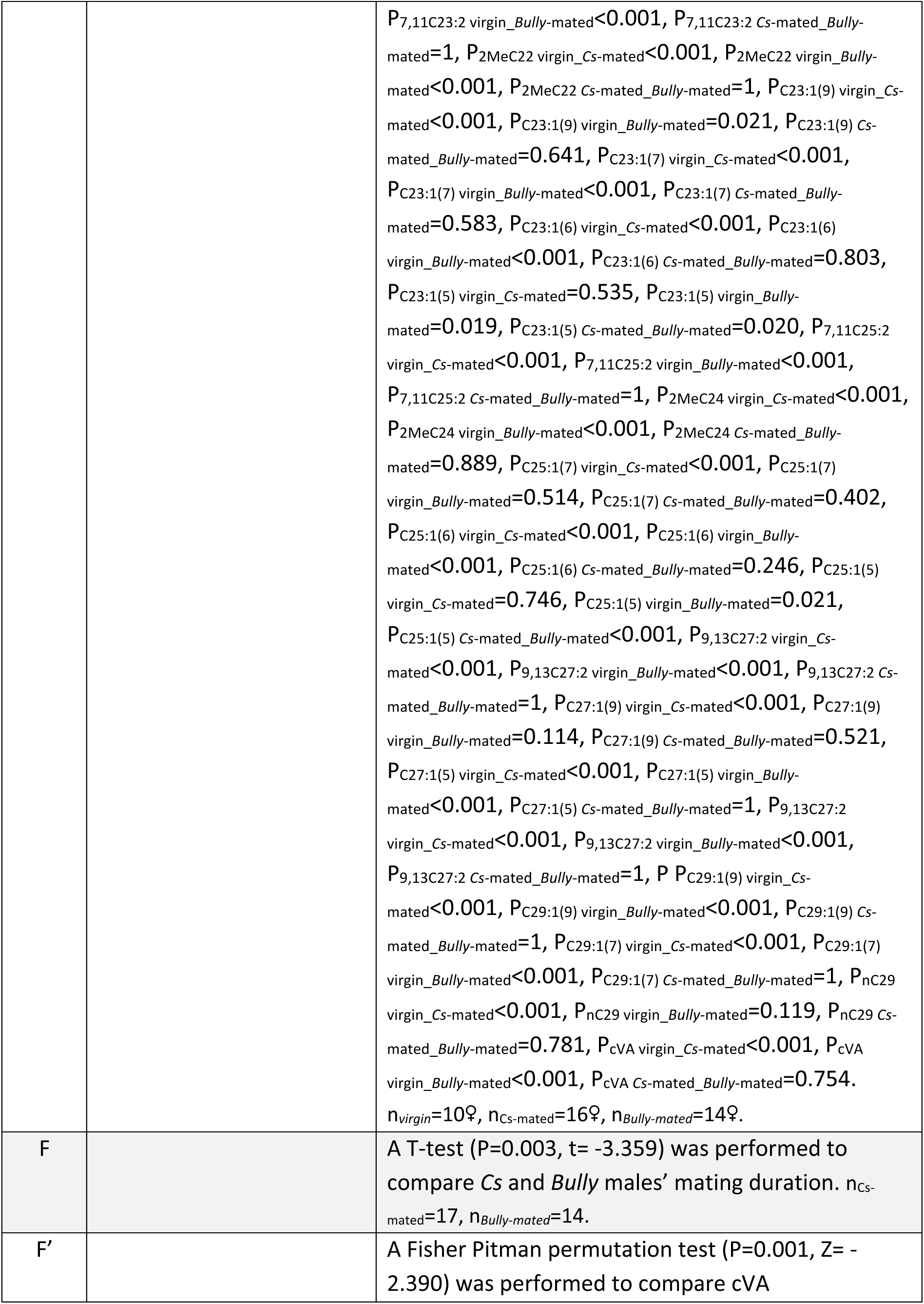
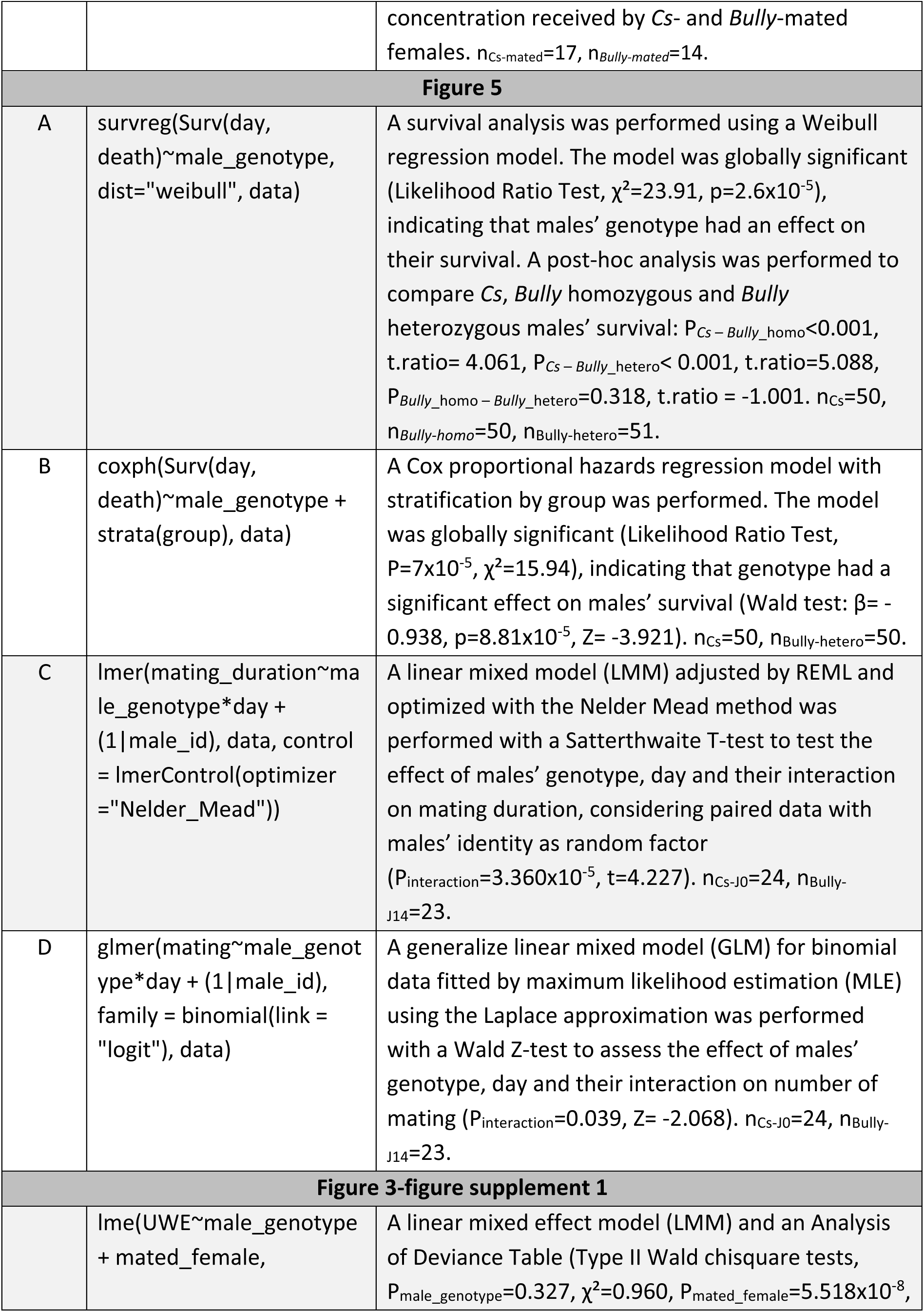
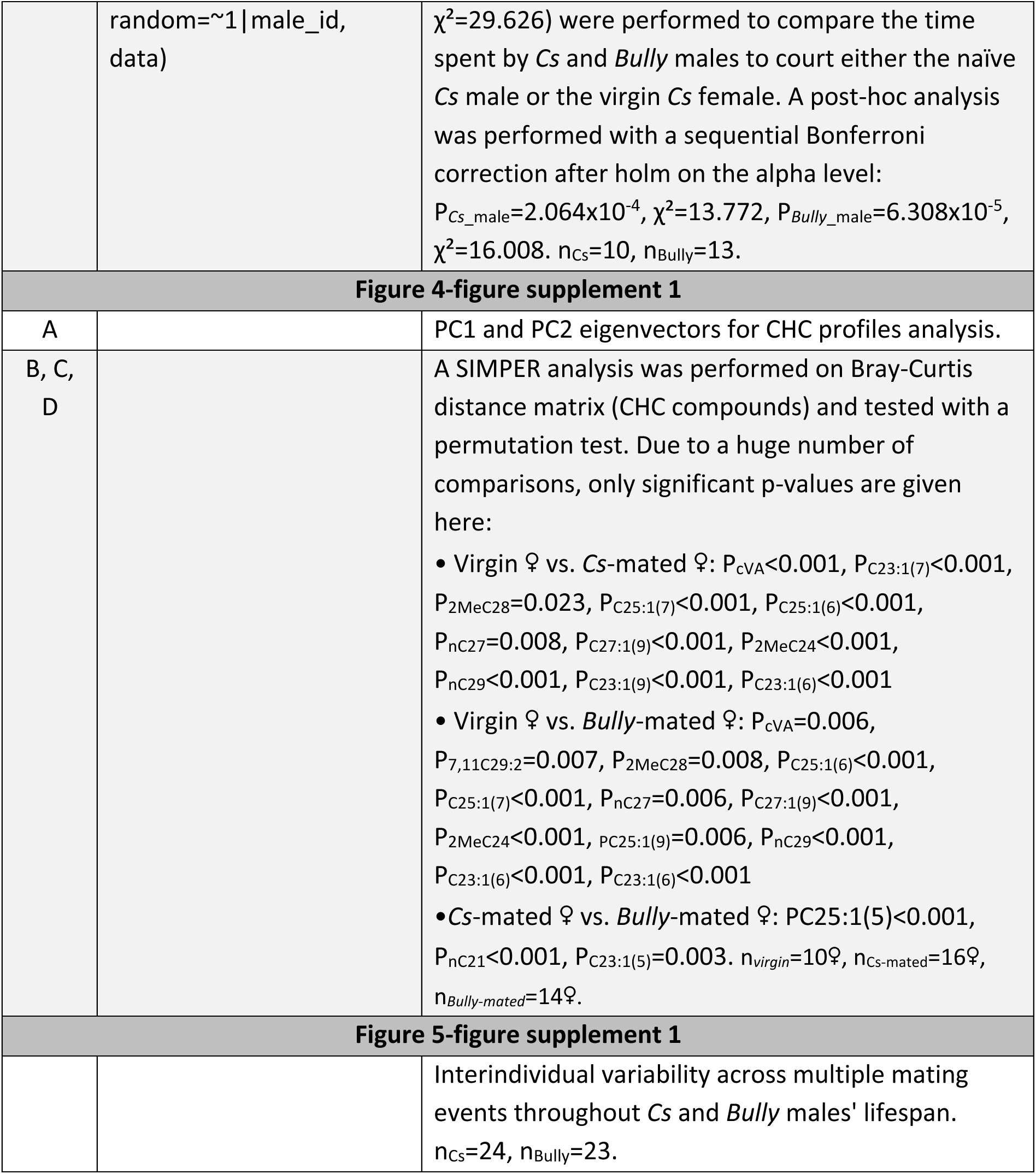
Statistical details of all analyses. This table gathers all statistical analyses performed in all the Figures. The first column refers to the panels of each Figure, the second one gives the statistical model used to analyze data and the last one provides the names of statistical tests and corrections performed as well as all p-values and statistical values.

**Figure 3-figure supplement 1.**
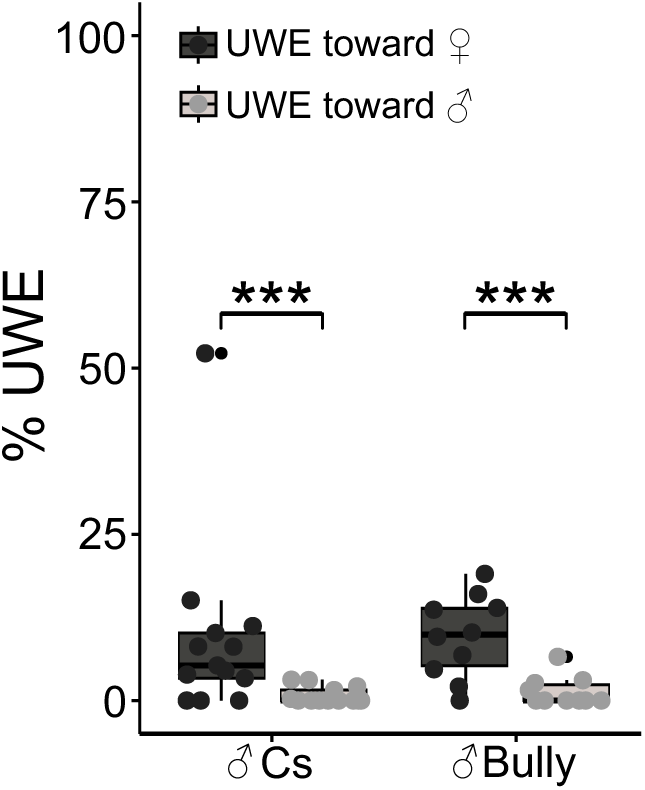
***Cs* and *Bully* males discrimination between males and females CHC profiles.** Time spent by *Cs* and *Bully* males to court either a naïve decapitated *Cs* male or a virgin *Cs* decapitated female (n_Cs_=10, n_Bully_=13, LMM and Analysis of Deviance Table (Type II Wald chisquare tests): P_male_genotype_=0.327, χ²=0.960, P_mated_female_=5.518×10^−8^, χ²=29.626). Stars represent significant differences (* P < 0.05, ** P < 0.01, *** P < 0.001). Post-hoc analyses and statistical details are given in Figure 1-table supplement 1.

**Figure 4-figure supplement 1.**
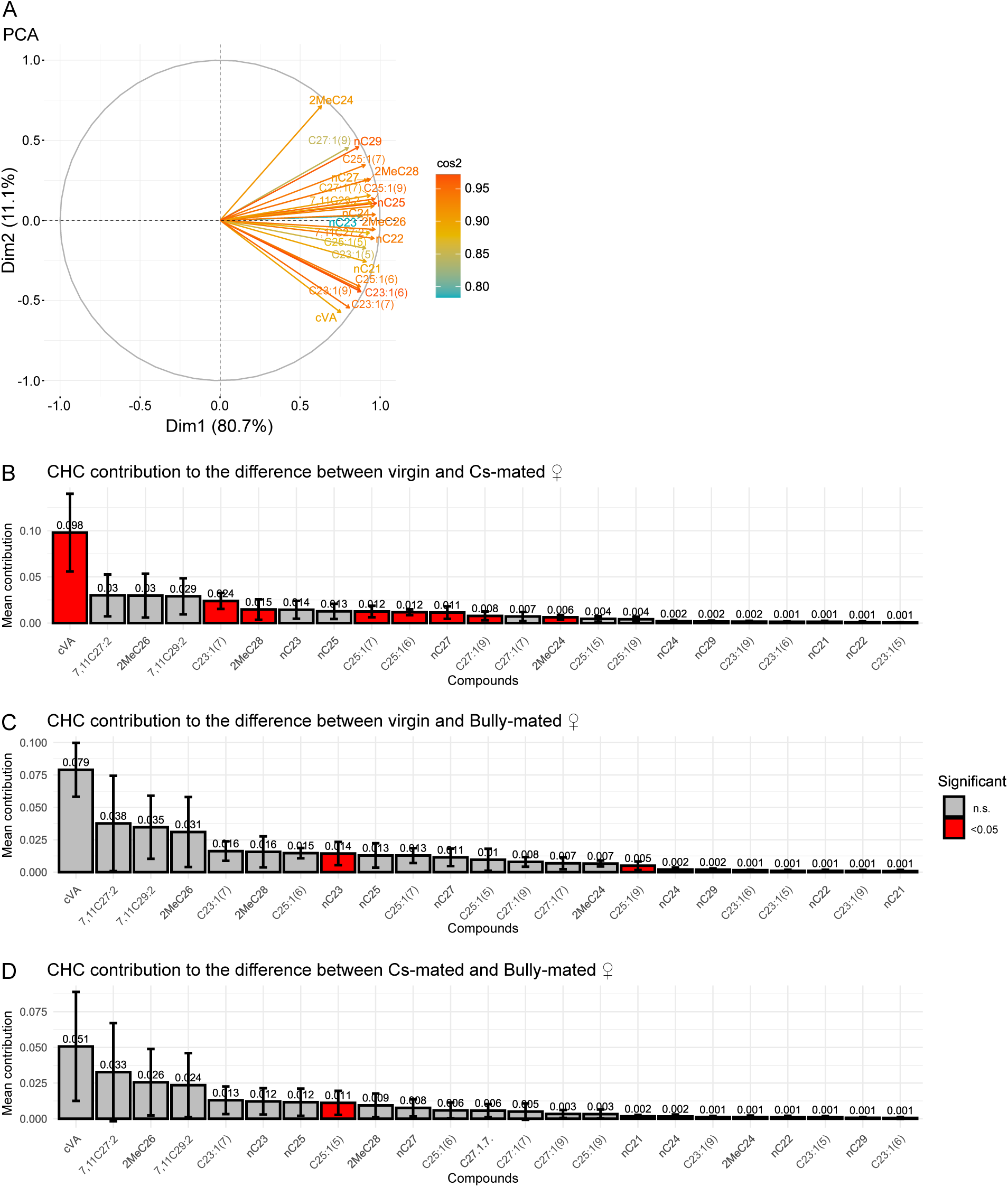
Details on PCA analysis for virgin and mated females. **(A)** Eigenvectors and associated cos² for each CHCs compound. Contribution of each CHCs compound in the difference between (**B)** virgin and *Cs*-mated, (**C)** virgin and *Bully*-mated and (**D)** *Cs*-mated and *Bully*-mated CHCs profiles. Compounds contributing significantly to the difference between CHCs profiles are represented in red. Numbers above bars refer to the mean contribution of each compound in the difference between CHCs profiles. Statistical details are given in Figure 1-table supplement 1. n*_virgin_*=10♀, n_Cs-mated_=16♀, n*_Bully-mated_*=14♀.

**Figure 5-figure supplement 1.**
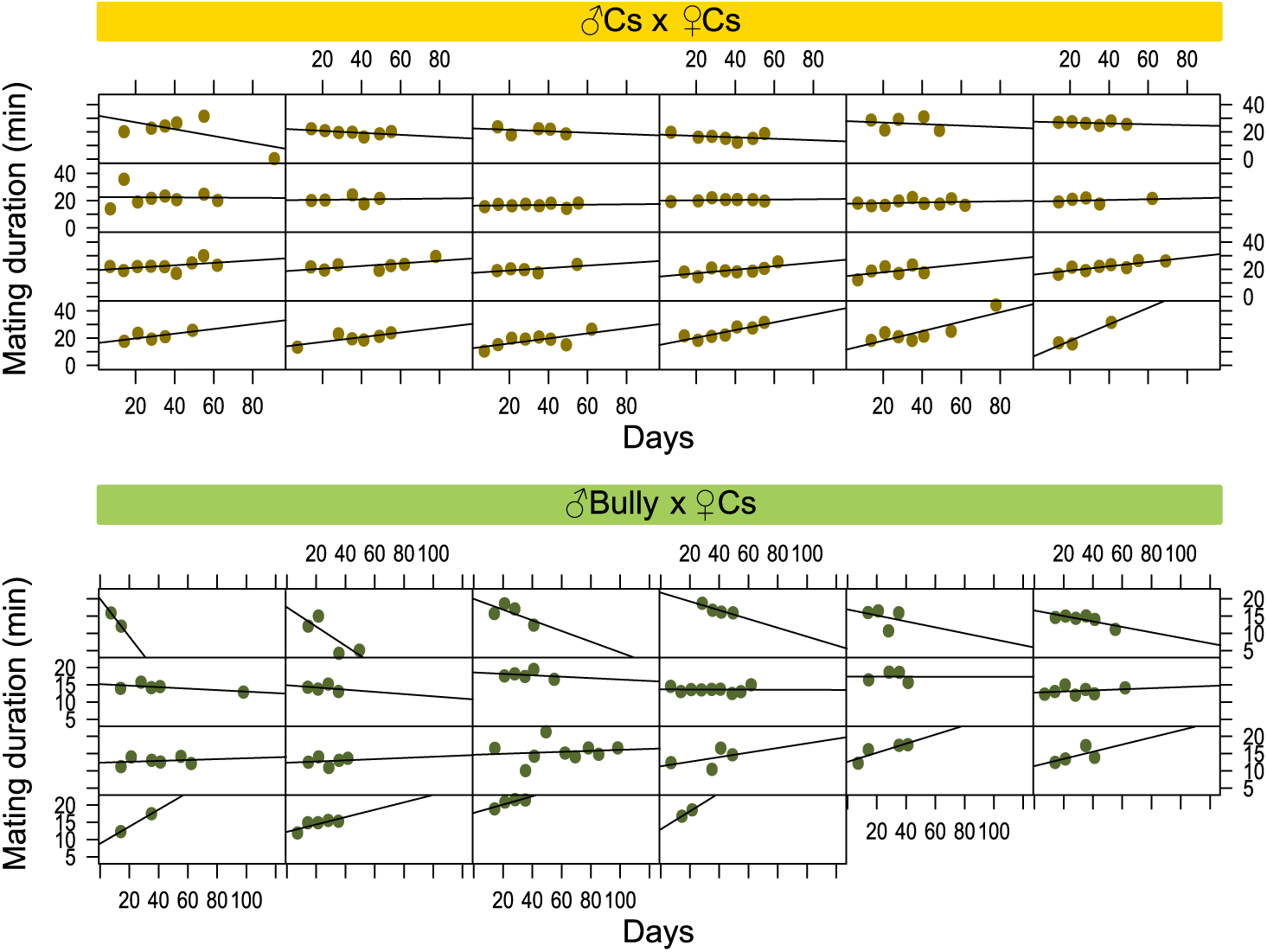
Interindividual variability across multiple mating events throughout the flies’ lifespan. Interindividual variability in mating duration for each (**A)** *Cs* and (**B)** *Bully* male when mated with a virgin *Cs* female across their lifespan. Each panel represents the mating durations (in minutes) for a single individual, with each dot corresponding to a single mating event. Statistical details are given in Figure 1-table supplement 1. n_Cs_=24, n_Bully_=23.

