## Supplementary material for "Selection for male aggression is associated with changes in reproductive traits, chemical signaling and lifespan in *Drosophila melanogaster*": Supp information

**SUPPORTING INFORMATIONS**

| Figure | Statistical model | Results |
|---|---|---|
| **Figure1** | | |
| B | lm(lunge_latency~males_genotype, data) | A linear model (LM) and a permutation test (Monte-Carlo test, P=0.136) was performed to compare *Cs*, *Bully* A, *Bully* B and *Bully* C males' latency to lunge. $n_{Cs}$=29, $n_{BullyA}$=24, $n_{BullyB}$=12, $n_{BullyC}$=11. |
| C | glm(nb_lunge~males_genotype, family=quasipoisson(link="log"), data) | A generalize linear model (GLM) for quasi-Poisson distributed data and a type II analysis of deviance table (Likelihood Ratio Test, P=1.622x$10^{-10}$, LR$\chi^2$=48.556) were performed to compare *Cs*, *Bully* A, *Bully* B and *Bully* C males' number of lunges. A post-hoc analysis was performed with a sequential Bonferroni correction after Holm on the p-values: $P_{Cs\_BullyA}$<0.001, z ratio= -4.495, $P_{Cs\_BullyB}$<0.001, z.ratio= -5.911, $P_{Cs\_BullyC}$=0.018, z.ratio= -2.836, $P_{BullyA\_BullyB}$=0.042, z ratio= -2.308, $P_{BullyA\_BullyC}$=0.232, z ratio= 1.195, $P_{BullyB\_BullyC}$=0.018, z ratio=2.797. $n_{Cs}$=29, $n_{BullyA}$=24, $n_{BullyB}$=12, $n_{BullyC}$=11. |
| D | glm(nb_boxe~genotype, family=poisson(link="log"), data) | A generalize linear model (GLM) for Poisson distributed data and a type II analysis of deviance table (Likelihood Ratio Test, P=2.2x$10^{-16}$, LR$\chi^2$=2511.9) were performed to compare *Cs*, *Bully* A, *Bully* B and *Bully* C males' number of boxing events. A post-hoc analysis was performed with a sequential Bonferroni correction after Holm on the p-values: $P_{Cs\_BullyA}$<0.001, z ratio= -6.819, $P_{Cs\_BullyB}$<0.001, z.ratio= -6.839, $P_{Cs\_BullyC}$<0.001, z.ratio= -3.649, $P_{BullyA\_BullyB}$=0.728, z ratio= -0.347, $P_{BullyA\_BullyC}$<0.001, z ratio=5.392, $P_{BullyB\_BullyC}$<0.001, z ratio=5.366. $n_{Cs}$=29, $n_{BullyA}$=24, $n_{BullyB}$=12, $n_{BullyC}$=11. |
| E | glm(mating_success~genotype, family=binomial(link= "logit"), data) | A generalize linear model (GLM) for binomial data and a type II analysis of deviance table (Likelihood Ratio Test, P=0.012, LR$\chi^2$=11.040) were performed to compare *Cs*, *Bully* A, *Bully* B and *Bully* C males' mating success. A post-hoc analysis was performed with a sequential Bonferroni correction after Holm on the p-values: $P_{Cs\_BullyA}$=0.934, z ratio=0.728, $P_{Cs\_BullyB}$=0.099, z.ratio=2.395, $P_{Cs\_BullyC}$=0.868, z.ratio=1.060, $P_{BullyA\_BullyB}$=0.206, z |

| | | |
|---|---|---|
| | | ratio=2.043, $P_{BullyA\_BullyC}$=0.934, z ratio=0.406, $P_{BullyB\_BullyC}$=0.522, z ratio= -1.513. P-values were not significant after correction. $n_{Cs}$=37, $n_{BullyA}$=31, $n_{BullyB}$=33, $n_{BullyC}$=21. |
| F | lm(UWE~males_genotype, data) | A linear model (LM) and a type II Anova table (P=0.088, F=2.235) were performed to compare *Cs*, *Bully* A, *Bully* B and *Bully* C males' time spent to court the virgin female. $n_{Cs}$=37, $n_{BullyA}$=31, $n_{BullyB}$=33, $n_{BullyC}$=21. |
| G | lm(courtship_latency~males_genotype, data) | A linear model (LM) and a type II Anova table (P=0.014, F=3.707) were performed to compare *Cs*, *Bully* A, *Bully* B and *Bully* C males' latency to court. A post-hoc analysis was performed with a sequential Bonferroni correction after Holm on the p-values: $P_{Cs\_BullyA}$=0.338, t.ratio= -1.740, $P_{Cs\_BullyB}$=0.433, t.ratio= 1.469, $P_{Cs\_BullyC}$=0.524, t.ratio= -1.127, $P_{BullyA\_BullyB}$=0.015, t.ratio= 3.101, $P_{BullyA\_BullyC}$=0.683, t.ratio=0.661, $P_{BullyB\_BullyC}$=0.099, t.ratio= -2.364. $n_{Cs}$=37, $n_{BullyA}$=31, $n_{BullyB}$=33, $n_{BullyC}$=21. |
| H | lm(mating_latency~males_genotype, data) | A linear model (LM) and a type II Anova table ($P=8.527x10^{-4}$, F=5.970) were performed to compare *Cs*, *Bully* A, *Bully* B and *Bully* C males' latency to mate. A post-hoc analysis was performed with a sequential Bonferroni correction after Holm on the p-values: $P_{Cs\_BullyA}$=0.482, t.ratio= -1.413, $P_{Cs\_BullyB}$<0.001, t.ratio= -4.191, $P_{Cs\_BullyC}$=0.581, t.ratio= -1.062, $P_{BullyA\_BullyB}$=0.038, t.ratio= -2.725, $P_{BullyA\_BullyC}$=0.863, t.ratio=0.173, $P_{BullyB\_BullyC}$=0.041, t.ratio= 2.615. $n_{Cs}$=36, $n_{BullyA}$=29, $n_{BullyB}$=24, $n_{BullyC}$=19. |
| I | lm(mating_duration~males_genotype, data) | A linear model (LM) and a type II Anova table ($P=5.015x10^{-6}$, F=10.365) were performed to compare *Cs*, *Bully* A, *Bully* B and *Bully* C males' mating duration. A post-hoc analysis was performed with a sequential Bonferroni correction after Holm on the p-values: $P_{Cs\_BullyA}$<0.001, t.ratio=4.537, $P_{Cs\_BullyB}$<0.001, t.ratio= 4.849, $P_{Cs\_BullyC}$=0.087, t.ratio= 2.332, $P_{BullyA\_BullyB}$=0.599, t.ratio=0.528, $P_{BullyA\_BullyC}$=0.227, t.ratio= -1.595, $P_{BullyB\_BullyC}$=0.142, t.ratio= -2.008. $n_{Cs}$=36, $n_{BullyA}$=29, $n_{BullyB}$=24, $n_{BullyC}$=19 |
| **Figure 2** | | |
| A | glm(mating~male_genotype, family=binomial(link="logit"), data) | A generalize linear model (GLM) for binomial data and a type II analysis of deviance table (Likelihood Ratio Test, |

| | | |
|---|---|---|
| | | P=0.329, LR$\chi^2$=0.951) were performed to compare *Cs* and *Bully* males' mating success. $n_{Cs}$=19, $n_{Bully}$=19. |
| B | glm(mating~male_genotype, family=binomial(link="logit"), data) | A generalize linear model (GLM) for binomial data and a type II analysis of deviance table (Likelihood Ratio Test, P=0.037, LR$\chi^2$=4.347) were performed to compare *Cs* and *Bully* males' mating success. $n_{Cs}$=64, $n_{Bully}$=41. |
| C | | A Fisher Pitman permutation test (P=0.877, t= -0.131) was performed to compare the proportion of time spent by *Cs* and *Bully* males to court the virgin female. $n_{Cs}$=64, $n_{Bully}$=41. |
| D | | A Fisher Pitman permutation test (P=0.319, Z=1.043) was performed to compare *Cs* and *Bully* males' latency to court. $n_{Cs}$=64, $n_{Bully}$=41. |
| E | | A Fisher Pitman permutation test (P=0.266, Z=1.170) was performed to compare *Cs* and *Bully* males' latency to mate. $n_{Cs}$=61, $n_{Bully}$=34. |
| F | | A T-test (P=6.009x10$^{-10}$, t= -6.917) was performed to compare *Cs* and *Bully* males' mating duration. $n_{Cs}$=61, $n_{Bully}$=34. |
| G | | A T-test (P=0.022, t= -2.367) was performed to compare *Cs* and *Bully* males' mating duration. $n_{Cs}$=61, $n_{Bully}$=34. $n_{Cs}$=26, $n_{Bully}$=20. |
| H | lm(mating_duration~ condition, data) | A linear model (LM) and a type II ANOVA (F-test, P=7.582x10$^{-11}$, F=21.669) were performed to compare *Cs* and *Bully* males' mating duration among raising condition. A post-hoc analysis was performed with a sequential Bonferroni correction after Holm on the alpha level: $P_{Cs\text{-isolated\_}Cs\text{-group}}$=0.013, F=6.583, $P_{Bully\text{-isolated\_}Bully\text{-group}}$=2.532x10$^{-4}$, F=15.731, $P_{Cs\text{-isolated\_}Bully\text{-isolated}}$=4.263x10$^{-6}$, F=27.697, $P_{Cs\text{-group\_}Bully\text{-group}}$=1.884x10$^{-4}$, F=16.032. $n_{Cs\ isolated}$=22, $n_{Cs\ group}$=32, $n_{Bully\ isolated}$=23, $n_{Bully\ group}$=25. |
| **Figure 3** | | |
| A | | A T-test (P=2.912x10$^{-11}$, t= -8.759) was performed to compare *Cs* and *Bully* males' mating duration. $n_{Cs}$=21, $n_{Bully}$=22. |
| A' | | A Fisher Pitman permutation test (P<0.001, Z=3.239) was performed to compare *Cs* and *Bully* males' time spent to court the mated female. $n_{Cs}$=21, $n_{Bully}$=22. |
| B | | A T-test (P=5.425x10$^{-7}$, t= -5.818) was performed to compare *Cs* and *Bully* males' mating duration. $n_{Cs}$=24, $n_{Bully}$=24. |

| | | |
|---|---|---|
| B’ | | A Fisher Pitman permutation test ($P=0.007$, $Z= -2.679$) was performed to compare *Cs* and *Bully* males’ time spent to court the mated female. $n_{Cs}=24$, $n_{Bully}=24$. |
| C | glmer(mating~male_genotype + (1 \| male_id), family = binomial(link = "logit"), data) | A type II analysis of deviance table (Wald Chi-square test, $P=0.659$, $\chi^2=0.195$) was performed to compare *Cs* and *Bully* males’ total number of mating. |
| D | lmer(mating_duration~ couples*mating + (1\|male_id), data) | A linear mixed model (LMM) adjusted by REML and a Satterthwaite T-test were performed to test the effect of males’ genotype, number of mating and their interaction on mating duration, considering paired data with males’ identity as random factor ($P_{interaction}=0.001$, $t= -3.261$). |
| E | glmer(progeny~male_genotype + mating + (1\|male_id), family=binomial(link="logit"), data) | A generalized linear mixed model (GLMM) and a Wald Chi-square test were performed to test the effect of males’ genotype and number of mating on number of mating giving progeny, considering paired data with males’ identity as random factor ($P_{male\_genotype}=0.279$, $\chi^2=1.173$, $P_{mating}=0.847$, $\chi^2=2.019$). |
| F | | A T-test ($P=7.139\times10^{-4}$, $t= -3.617$) was performed to compare *Cs* and *Bully* males’ mating duration. $n_{Cs}=25$, $n_{Bully}=25$. |
| F’ | lme(UWE~male_genotype + mated_female, random=~1\|male_id, data) | A linear mixed effect model (LMM) and a permutation test (Monte-Carlo test, $P_{male\_genotype}<0.001$, $P_{mated\_female}=0.457$) was performed to compare *Cs* and *Bully* males’ time spent to court the mated female and if *Cs*- and *Bully*-mated females received the same amount of courtship. $n_{Cs\text{-}Cs\text{-}mated}=12$, $n_{Cs\text{-}Bully\text{-}mated}=15$, $n_{Bully\text{-}Cs\text{-}mated}=9$, $n_{Bully\text{-}Bully\text{-}mated}=10$. |
| G | | A T-test ($P=8.743\times10^{-5}$, $t= -4.255$) was performed to compare *Cs* and *Bully* males’ mating duration. $n_{Cs}=30$, $n_{Bully}=24$. |
| G’ | glm(remating~mated_female + day, family = binomial(link = "logit"), data) | A generalize linear model (GLM) for binomial data and a type II analysis of deviance table (Likelihood Ratio Test, $P_{mated\_female}=3.639\times10^{-4}$, $LR\chi^2=12.708$, $P_{day}=0.385$, $LR\chi^2=0.756$) were performed to compare *Cs*- and *Bully*-mated re-mating proportion. $n_{Cs\text{-}12days}=19$, $n_{Bully\text{-}12days}=15$, $n_{Cs\text{-}14days}=11$, $n_{Bully\text{-}14days}=9$. |
| **Figure 4** | | |
| A | | A Principal Component Analysis (PCA) and a PerMANOVA were performed to test the effect of males’ genotype on Euclidian distances of individuals on CHC compounds ($P=0.002$, $F=6.789$, $R^2=0.274$). $n_{Cs}=10$♂, $n_{Bully}=10$♂. |

| | | |
|---|---|---|
| B + Table 1 | lme(concentration~male_genotype, random=~1\|male_id, data) | A linear mixed effect model (LMM) and a permutation test (Monte-Carlo test, P=0.009) were performed to compare CHC concentrations between naïve *Cs* and bully males. A post-hoc analysis was performed with a sequential Bonferroni correction after Benjamini and Hochberg on the alpha level: $P_{Alkanes}$=0.036, $P_{Monoenes}$=0.002, $P_{Methyl\ alkanes}$=0.048, $P_{nC21}$<0.001, $P_{C22:1(9)}$=0.892, $P_{C22:1(7)}$=0.568, $P_{cVA}$=0.731, $P_{nC22}$=0.854, $P_{C23:1(9)}$=0.175, $P_{C23:1(7)}$=0.062, $P_{C23:1(5)}$<0.001, $P_{nC23}$=0.014, $P_{C24:1(9)}$=0.345, $P_{C24:1(7}$<0.001, $P_{24:1(5)}$=0.015, $P_{nC24}$=0.002, $P_{2MeC24}$=0.548, $P_{C25:1(9)}$=0.524, $P_{C25:1(7)}$<0.001, $P_{C25:1(5)}$<0.001, $P_{nC25}$<0.001, $P_{2MeC26}$=0.159, $P_{C27:1(7)}$<0.001, $P_{nC27}$=0.078, $P_{2MeC28}$=0.003, $P_{nC29}$=0.005. $n_{\mathit{Cs}}$=10♂, $n_{\mathit{Bully}}$=10♂. |
| C | | A Principal Component Analysis (PCA) and a PerMANOVA were performed to test the effect of females' mating status on Euclidian distances of individuals on CHC compounds (P=0.017, F=2.594, $R^2$=0.123). A post-hoc analysis was performed with a sequential Bonferroni correction after holm on the alpha level: $P_{virgin\_\mathit{Cs}\text{-}mated}$=0.001, F=19.434, $R^2$=0.447, $P_{virgin\_\mathit{Bully}\text{-}mated}$=0.034, F=2.266, $R^2$=0.093, $P_{\mathit{Cs}\text{-}mated\_\mathit{Bully}\text{-}mated}$=0.746, F=0.428, $R^2$=0.015. $n_{\mathit{virgin}}$=10♀, $n_{Cs\text{-}mated}$=16♀, $n_{\mathit{Bully\text{-}mated}}$=14♀. |
| D + E + Table 2 | lme(concentration~female_status, random=~1\|female_id, data) | A linear mixed effect model (LMM) and a permutation test (Monte-Carlo test, P=0.618) were performed to compare CHC concentrations between virgin, *Cs*- and *Bully-mated* females. Although the global multivariate test did not detect a significant difference between females, we performed a family-wise comparison: $P_{Alkanes}$=0.705, $P_{Monoenes}$=0.294, $P_{Dienes}$=0.715, $P_{Methyl\ alkanes}$=0.750. The absence of a significant multivariate effect does not preclude compound-specific differences, as the multivariate tests evaluated the joint profile across all 28 compounds rather than the effect on any individual compound. Therefore, a compound-wise permutation analyses was performed: $P_{nC21}$=0.158, $Pn_{C22}$=0.927, $P_{7,11C23:2}$<0.001, $P_{2MeC22}$<0.001, $P_{C23:1(9)}$=0.023, $P_{C23:1(7)}$=0.001, $P_{C23:1(6)}$<0.001, $P_{C23:1(5)}$=0.016, $P_{nC23}$=0.908, $P_{nC24}$=0.899, $P_{7,11C25:2}$<0.001, $P_{2MeC24}$<0.001, $P_{C25:1(9)}$=0.832, $P_{C25:1(7)}$=0.047, $P_{C25:1(6)}$<0.001, $P_{C25:1(5)}$<0.001, $P_{nC25}$=0.769, $P_{9,13C27:2}$<0.001, $P_{7,11C27:2}$=0.747, $P_{2MeC26}$=0.884, |

| | | |
|---|---|---|
| | | $P_{\text{C27:1(9)}}=0.016$, $P_{\text{C27:1(7)}}=0.625$, $P_{\text{C27:1(5)}}<0.001$, $P_{\text{nC27}}=0.367$, $P_{\text{9,13C29:2}}<0.001$, $P_{\text{7,11C23:2}}=0.899$, $P_{\text{2MeC28}}=0.413$, $P_{\text{C29:1(9)}}<0.001$, $P_{\text{C29:1(7)}}<0.001$, $P_{\text{nC29}}=0.022$, $P_{\text{cVA}}=0.001$. For significant p-values, a post-hoc analysis was performed with a sequential Bonferroni correction after Holm on the alpha level to compare females within each compound: $P_{\text{7,11C23:2 virgin\_\textit{Cs}-mated}}<0.001$, $P_{\text{7,11C23:2 virgin\_\textit{Bully}-mated}}<0.001$, $P_{\text{7,11C23:2 \textit{Cs}-mated\_\textit{Bully}-mated}}=1$, $P_{\text{2MeC22 virgin\_\textit{Cs}-mated}}<0.001$, $P_{\text{2MeC22 virgin\_\textit{Bully}-mated}}<0.001$, $P_{\text{2MeC22 \textit{Cs}-mated\_\textit{Bully}-mated}}=1$, $P_{\text{C23:1(9) virgin\_\textit{Cs}-mated}}<0.001$, $P_{\text{C23:1(9) virgin\_\textit{Bully}-mated}}=0.021$, $P_{\text{C23:1(9) \textit{Cs}-mated\_\textit{Bully}-mated}}=0.641$, $P_{\text{C23:1(7) virgin\_\textit{Cs}-mated}}<0.001$, $P_{\text{C23:1(7) virgin\_\textit{Bully}-mated}}<0.001$, $P_{\text{C23:1(7) \textit{Cs}-mated\_\textit{Bully}-mated}}=0.583$, $P_{\text{C23:1(6) virgin\_\textit{Cs}-mated}}<0.001$, $P_{\text{C23:1(6) virgin\_\textit{Bully}-mated}}<0.001$, $P_{\text{C23:1(6) \textit{Cs}-mated\_\textit{Bully}-mated}}=0.803$, $P_{\text{C23:1(5) virgin\_\textit{Cs}-mated}}=0.535$, $P_{\text{C23:1(5) virgin\_\textit{Bully}-mated}}=0.019$, $P_{\text{C23:1(5) \textit{Cs}-mated\_\textit{Bully}-mated}}=0.020$, $P_{\text{7,11C25:2 virgin\_\textit{Cs}-mated}}<0.001$, $P_{\text{7,11C25:2 virgin\_\textit{Bully}-mated}}<0.001$, $P_{\text{7,11C25:2 \textit{Cs}-mated\_\textit{Bully}-mated}}=1$, $P_{\text{2MeC24 virgin\_\textit{Cs}-mated}}<0.001$, $P_{\text{2MeC24 virgin\_\textit{Bully}-mated}}<0.001$, $P_{\text{2MeC24 \textit{Cs}-mated\_\textit{Bully}-mated}}=0.889$, $P_{\text{C25:1(7) virgin\_\textit{Cs}-mated}}<0.001$, $P_{\text{C25:1(7) virgin\_\textit{Bully}-mated}}=0.514$, $P_{\text{C25:1(7) \textit{Cs}-mated\_\textit{Bully}-mated}}=0.402$, $P_{\text{C25:1(6) virgin\_\textit{Cs}-mated}}<0.001$, $P_{\text{C25:1(6) virgin\_\textit{Bully}-mated}}<0.001$, $P_{\text{C25:1(6) \textit{Cs}-mated\_\textit{Bully}-mated}}=0.246$, $P_{\text{C25:1(5) virgin\_\textit{Cs}-mated}}=0.746$, $P_{\text{C25:1(5) virgin\_\textit{Bully}-mated}}=0.021$, $P_{\text{C25:1(5) \textit{Cs}-mated\_\textit{Bully}-mated}}<0.001$, $P_{\text{9,13C27:2 virgin\_\textit{Cs}-mated}}<0.001$, $P_{\text{9,13C27:2 virgin\_\textit{Bully}-mated}}<0.001$, $P_{\text{9,13C27:2 \textit{Cs}-mated\_\textit{Bully}-mated}}=1$, $P_{\text{C27:1(9) virgin\_\textit{Cs}-mated}}<0.001$, $P_{\text{C27:1(9) virgin\_\textit{Bully}-mated}}=0.114$, $P_{\text{C27:1(9) \textit{Cs}-mated\_\textit{Bully}-mated}}=0.521$, $P_{\text{C27:1(5) virgin\_\textit{Cs}-mated}}<0.001$, $P_{\text{C27:1(5) virgin\_\textit{Bully}-mated}}<0.001$, $P_{\text{C27:1(5) \textit{Cs}-mated\_\textit{Bully}-mated}}=1$, $P_{\text{9,13C27:2 virgin\_\textit{Cs}-mated}}<0.001$, $P_{\text{9,13C27:2 virgin\_\textit{Bully}-mated}}<0.001$, $P_{\text{9,13C27:2 \textit{Cs}-mated\_\textit{Bully}-mated}}=1$, P $P_{\text{C29:1(9) virgin\_\textit{Cs}-mated}}<0.001$, $P_{\text{C29:1(9) virgin\_\textit{Bully}-mated}}<0.001$, $P_{\text{C29:1(9) \textit{Cs}-mated\_\textit{Bully}-mated}}=1$, $P_{\text{C29:1(7) virgin\_\textit{Cs}-mated}}<0.001$, $P_{\text{C29:1(7) virgin\_\textit{Bully}-mated}}<0.001$, $P_{\text{C29:1(7) \textit{Cs}-mated\_\textit{Bully}-mated}}=1$, $P_{\text{nC29 virgin\_\textit{Cs}-mated}}<0.001$, $P_{\text{nC29 virgin\_\textit{Bully}-mated}}=0.119$, $P_{\text{nC29 \textit{Cs}-mated\_\textit{Bully}-mated}}=0.781$, $P_{\text{cVA virgin\_\textit{Cs}-mated}}<0.001$, $P_{\text{cVA virgin\_\textit{Bully}-mated}}<0.001$, $P_{\text{cVA \textit{Cs}-mated\_\textit{Bully}-mated}}=0.754$. $n_{\textit{virgin}}=10$♀, $n_{\text{Cs-mated}}=16$♀, $n_{\textit{Bully-mated}}=14$♀. |
| F | | A T-test (P=0.003, t= -3.359) was performed to compare *Cs* and *Bully* males' mating duration. $n_{\text{Cs-mated}}=17$, $n_{\textit{Bully-mated}}=14$. |
| F' | | A Fisher Pitman permutation test (P=0.001, Z= -2.390) was performed to compare cVA concentration received |

| | | |
|---|---|---|
| | | by *Cs*- and *Bully*-mated females. $n_{Cs\text{-}mated}$=17, $n_{Bully\text{-}mated}$=14. |
| **Figure 5** | | |
| A | survreg(Surv(day, death)~male_genotype, dist="weibull", data) | A survival analysis was performed using a Weibull regression model. The model was globally significant (Likelihood Ratio Test, $\chi^2$=23.91, p=2.6x$10^{-5}$), indicating that males' genotype had an effect on their survival. A post-hoc analysis was performed to compare *Cs*, *Bully* homozygous and *Bully* heterozygous males' survival: $P_{Cs - Bully\_homo}$<0.001, t.ratio= 4.061, $P_{Cs - Bully\_hetero}$< 0.001, t.ratio=5.088, $P_{Bully\_homo - Bully\_hetero}$=0.318, t.ratio = -1.001. $n_{Cs}$=50, $n_{Bully\text{-}homo}$=50, $n_{Bully\text{-}hetero}$=51. |
| B | coxph(Surv(day, death)~male_genotype + strata(group), data) | A Cox proportional hazards regression model with stratification by group was performed. The model was globally significant (Likelihood Ratio Test, P=7x$10^{-5}$, $\chi^2$=15.94), indicating that genotype had a significant effect on males' survival (Wald test: β= -0.938, p=8.81x$10^{-5}$, Z= -3.921). $n_{Cs}$=50, $n_{Bully\text{-}hetero}$=50. |
| C | lmer(mating_duration~male_genotype*day + (1\|male_id), data, control = lmerControl(optimizer ="Nelder_Mead")) | A linear mixed model (LMM) adjusted by REML and optimized with the Nelder Mead method was performed with a Satterthwaite T-test to test the effect of males' genotype, day and their interaction on mating duration, considering paired data with males' identity as random factor ($P_{interaction}$=3.360x$10^{-5}$, t=4.227). $n_{Cs\text{-}J0}$=24, $n_{Bully\text{-}J14}$=23. |
| D | glmer(mating~male_genotype*day + (1\|male_id), family = binomial(link = "logit"), data) | A generalize linear mixed model (GLM) for binomial data fitted by maximum likelihood estimation (MLE) using the Laplace approximation was performed with a Wald Z-test to assess the effect of males' genotype, day and their interaction on number of mating ($P_{interaction}$=0.039, Z= -2.068). $n_{Cs\text{-}J0}$=24, $n_{Bully\text{-}J14}$=23. |
| **Figure 3-figure supplement 1** | | |
| | lme(UWE~male_genotype + mated_female, random=~1\|male_id, data) | A linear mixed effect model (LMM) and an Analysis of Deviance Table (Type II Wald chisquare tests, $P_{male\_genotype}$=0.327, $\chi^2$=0.960, $P_{mated\_female}$=5.518x$10^{-8}$, $\chi^2$=29.626) were performed to compare the time spent by *Cs* and *Bully* males to court either the naïve *Cs* male or the virgin *Cs* female. A post-hoc analysis was performed with a sequential Bonferroni correction after holm on the alpha level: $P_{Cs\_male}$=2.064x$10^{-4}$, $\chi^2$=13.772, $P_{Bully\_male}$=6.308x$10^{-5}$, $\chi^2$=16.008. $n_{Cs}$=10, $n_{Bully}$=13. |

| Figure 4-figure supplement 1 | | |
|---|---|---|
| A | | PC1 and PC2 eigenvectors for CHC profiles analysis. |
| B, C, D | | A SIMPER analysis was performed on Bray-Curtis distance matrix (CHC compounds) and tested with a permutation test. Due to a huge number of comparisons, only significant p-values are given here:<br>• Virgin ♀ vs. *Cs*-mated ♀: $P_{cVA}$<0.001, $P_{C23:1(7)}$<0.001, $P_{2MeC28}$=0.023, $P_{C25:1(7)}$<0.001, $P_{C25:1(6)}$<0.001, $P_{nC27}$=0.008, $P_{C27:1(9)}$<0.001, $P_{2MeC24}$<0.001, $P_{nC29}$<0.001, $P_{C23:1(9)}$<0.001, $P_{C23:1(6)}$<0.001<br>• Virgin ♀ vs. *Bully*-mated ♀: $P_{cVA}$=0.006, $P_{7,11C29:2}$=0.007, $P_{2MeC28}$=0.008, $P_{C25:1(6)}$<0.001, $P_{C25:1(7)}$<0.001, $P_{nC27}$=0.006, $P_{C27:1(9)}$<0.001, $P_{2MeC24}$<0.001, $_{PC25:1(9)}$=0.006, $P_{nC29}$<0.001, $P_{C23:1(6)}$<0.001, $P_{C23:1(6)}$<0.001<br>•*Cs*-mated ♀ vs. *Bully*-mated ♀: PC25:1(5)<0.001, $P_{nC21}$<0.001, $P_{C23:1(5)}$=0.003. $n_{virgin}$=10♀, $n_{Cs\text{-}mated}$=16♀, $n_{Bully\text{-}mated}$=14♀. |
| **Figure 5-figure supplement 1** | | |
| | | Interindividual variability across multiple mating events throughout *Cs* and *Bully* males' lifespan. $n_{Cs}$=24, $n_{Bully}$=23. |

**Figure 1-table supplement 1. Statistical details of all analyses.** This table gathers all statistical analyses performed in all the Figures. The first column refers to the panels of each Figure, the second one gives the statistical model used to analyze data and the last one provides the names of statistical tests and corrections performed as well as all p-values and statistical values.

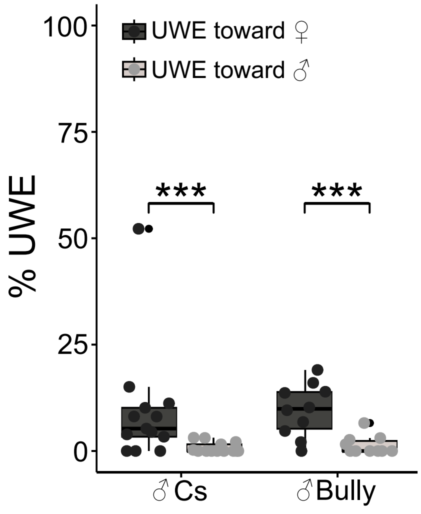


**Figure 3-figure supplement 1. *Cs* and *Bully* males discrimination between males and females CHC profiles.**

Time spent by *Cs* and *Bully* males to court either a naïve decapitated *Cs* male or a virgin *Cs* decapitated female ($n_{Cs}$=10, $n_{Bully}$=13, LMM and Analysis of Deviance Table (Type II Wald chisquare tests): $P_{male\_genotype}$=0.327, $\chi^2$=0.960, $P_{mated\_female}$=5.518x$10^{-8}$, $\chi^2$=29.626). Stars represent significant differences (* P < 0.05, ** P < 0.01, *** P < 0.001). Post-hoc analyses and statistical details are given in Figure 1-table supplement 1.

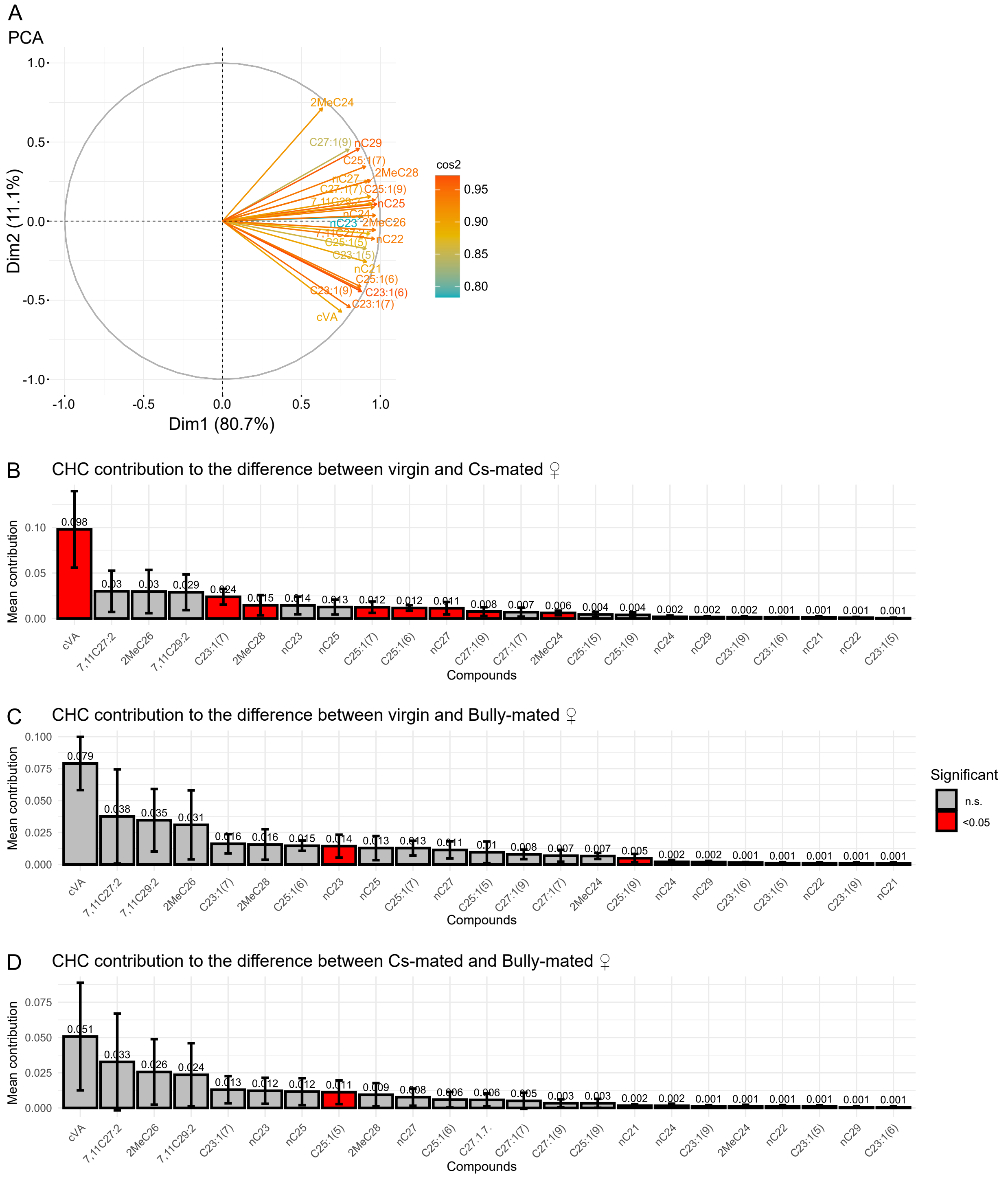


**Figure 4-figure supplement 1. Details on PCA analysis for virgin and mated females.**

**(A)** Eigenvectors and associated cos² for each CHCs compound. Contribution of each CHCs compound in the difference between **(B)** virgin and *Cs*-mated, **(C)** virgin and *Bully*-mated and **(D)** *Cs*-mated and *Bully*-mated CHCs profiles. Compounds contributing significantly to the difference between CHCs profiles are represented in red. Numbers above bars refer to the mean contribution of each compound in the difference between CHCs profiles. Statistical details are given in Figure 1-table supplement 1. $n_{virgin}$=10♀, $n_{Cs\text{-}mated}$=16♀, $n_{Bully\text{-}mated}$=14♀.

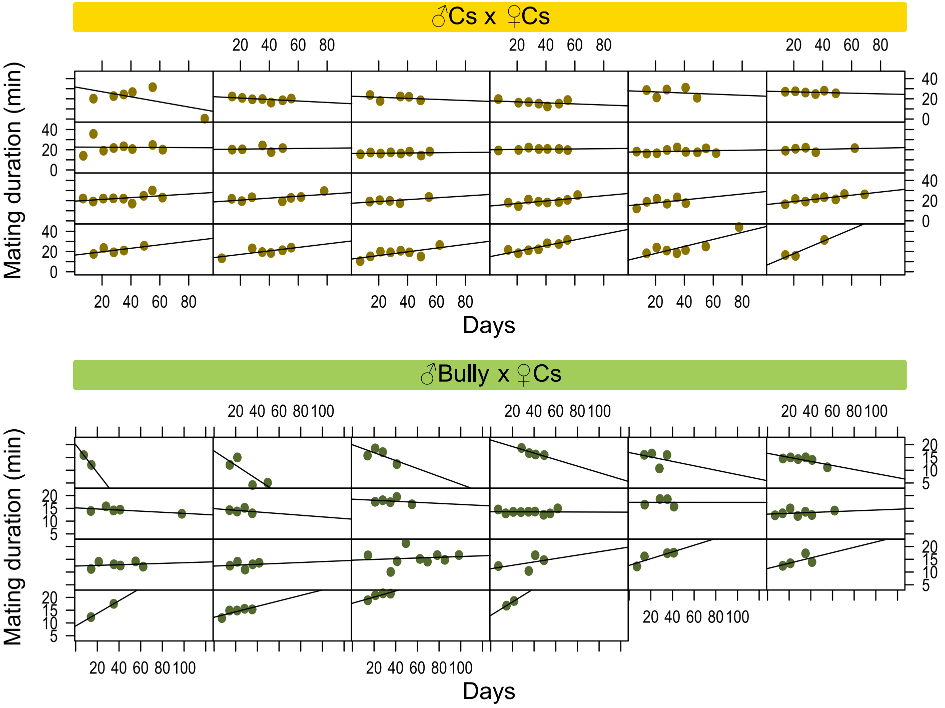


**Figure 5-figure supplement 1. Interindividual variability across multiple mating events throughout the flies' lifespan.** Interindividual variability in mating duration for each **(A)** *Cs* and **(B)** *Bully* male when mated with a virgin *Cs* female across their lifespan. Each panel represents the mating durations (in minutes) for a single individual, with each dot corresponding to a single mating event. Statistical details are given in Figure 1-table supplement 1. $n_{Cs}$=24, $n_{Bully}$=23.
